# Qualitative and Quantitative Comparative Analysis of Common Normal Variants and Physiological Artifacts in MEG and EEG

**DOI:** 10.1101/2025.03.24.645100

**Authors:** Daria Kleeva, Mikhail Sinkin, Anna Shtekleyn, Anna Rusinova, Anastasia Skalnaya, Alexei Ossadtchi

## Abstract

Magnetoencephalography (MEG) and electroencephalography (EEG) provide complementary insights into brain activity, yet their distinct biophysical principles influence how normal neurophysiological patterns and artifacts are represented. This study presents a comprehensive qualitative and quantitative analysis of common physiological variants and artifacts in simultaneously recorded MEG and EEG data. We systematically examined patterns such as alpha spindles, sensorimotor rhythms, sleep-related waveforms (vertex waves, K-complexes, sleep spindles, and posterior slow waves of youth), as well as common artifacts including eye blinks, chewing, and movement-related interferences. By applying time-domain, time-frequency, and source-space analyses, we identified modality-specific differences in signal representation, source localization, and artifact susceptibility. Our results demonstrate that MEG provides a more spatially focal representation of physiological patterns, whereas EEG captures broader, radially oriented cortical activity. Mutual information analysis indicated that MEG-derived independent components exhibited greater topographical variability and higher information content for neurophysiological activity, while EEG components were more homogeneous. Signal-to-noise ratio (SNR) analysis confirmed that MEG gradiometers capture the highest total information, followed by magnetometers and then EEG. Notably, physiological signals such as vertex waves and K-complexes exhibited significantly higher total information in MEG, whereas EEG was more sensitive to high-amplitude artifacts, including swallowing and muscle activity. These findings highlight the distinct strengths and limitations of MEG and EEG, reinforcing the necessity of multimodal approaches in clinical and research applications to improve the accuracy of neurophysiological assessments.

## Introduction

The human brain’s complex electrical activity remains a subject of immense interest and investigation in neuroscience. The assessment of cerebral bioelectrical activity forms the cornerstone of understanding brain function and diagnosing neurological disorders, such as depression and anxiety, Parkinson’s and Alzheimer’s diseases, epilepsy (Fred et al., 2022a; Zhang et al., 2022; Rassoulou et al., 2024; Schoonhoven et al., 2022; Laohathai et al., 2021). Magnetoencephalography (MEG) and Electroencephalography (EEG), while distinct in their methodologies, jointly pioneer this exploration (Fred et al., 2022b; Beniczky and Schomer, 2020). When applied to epilepsy the goal is to discover specific morphological patterns of brain’s electrical activity that hallmark pathological cortical zones.

Magneto- and electroencephalography (MEG and EEG) furnish non-invasive and time-resolved access to the brain’s function. EEG, by registering the electrical potentials generated by neural assemblies, provides a straightforward window into the electrical communication within the brain. On the other hand, magnetic fields produced by these electrical currents and registered with MEG enjoy greater informativeness (Hari and Puce, 2023) and allow for the improved spatial resolution of the cortical maps obtained with mathematical methods for solving the inverse problem King and Wyart (2021); Mosher et al. (1999) and detecting functional networks Brookes et al. (2011) including those coupled at zero or close to zero lag Ossadtchi et al. (2018).

Many MEG systems, such as the Elekta Neuromag, utilize two different sensor types– magnetometers and gradiometers – to capture magnetic signals associated with neural activity in a different manner. Magnetometers are sensitive to both superficial and deep cortical sources but their measurements are affected by the environmental interference. Gradiometers, on the other hand, designed to measure the spatial gradient of the magnetic field, are less sensitive to the external noises but are biased towards superficial sources. Combining these two types sensors allows for more informative measurements (Garćes et al., 2017) than it could be afforded by either of them. The outlined differences underscore the complementary nature of MEG and EEG in brain research and motivate their simultaneous usage which is technically possible Yoshinaga et al. (2002).

MEG’s forward model is less dependent upon the unknown distribution of conductivity with the head which makes it superior with respect to EEG in the source localization accuracy. At the same time, MEG is significantly less informative with respected to radially oriented sources than EEG which justifies the concurrent use of these two methods (Malmivuo, 2012). EEG and MEG may differently perceive the geometric details of localized neuronal activity (Lopes da Silva et al., 1991) and the ultimate source localization accuracy critically depends on the fidelity of the corresponding forward models. Under low signal-to-noise conditions, EEG source localization has been found to be more sensitive to head modeling errors as compared to MEG, which may limit EEG’s precision in noisy conditions (Cohen and Cuffin, 1991).

The use of high density EEG benefits source localization accuracy as compared to the low density EEG setups and makes the discovered sources concordant with anatomical references obtained by fMRI (Klamer et al., 2015).

Somewhat controversial observation was reported in (Liu et al., 2002) where the authors found that given the same number of sensors EEG is more accurate than MEG. MEG was found to have higher crosstalk and less focal point spread function, particularly for deep sources. The combination of EEG and MEG provided the best localization accuracy, resulted in the reduced crosstalk and more selective point spread functions than either of the two modalities alone. The other study performed on somatosensory evoked responses (Hedrich et al., 2017) compared several source imaging methods and demonstrated that MEG data exhibited better performance compared to hdEEG in terms of spatial resolution, largely due to MEG’s higher signal-to-noise ratio. Finally, MEG measurements interpreted with complex machine learning methods appear to be significantly more informative than those obtained with EEG during the speech comprehension task (Défossez et al., 2023).

Due to inherent differences in signal acquisition between MEG and EEG, distinct analytical frameworks are required for each. Specifically, MEG’s lack of a physical reference electrode, a cornerstone in EEG signal interpretation, renders traditional concepts, e.g. signal ”polarity”, harder to interpret. Additionally, MEG’s excessive channel count poses challenges for swift visual assessments, complicating the direct application of EEG’s clinical visual analysis habits possessed by clinicians. Moreover, as highlighted in Rampp et al. (2020), MEG sensor design affects the appearance of normal variants, with planar gradiometers and magnetometers showing different sensitivity to source depth and noise. This technical aspect reinforces the necessity of understanding the intricacies of MEG data analysis and the importance of simultaneous EEG recording to aid in the correct identification of non-pathological morphological patterns of brain activity. Furthermore, these differences also exacerbate the challenges faced by clinical neurophysiologists primarily experienced in making clinical decisions based on EEG data. The shift to MEG analysis requires not only a technical understanding of its unique data characteristics but also an adaptation of clinical decision-making processes, underscoring the need for specialized training and experience in MEG for those traditionally versed in EEG.

To aid this process, here we have implemented a comprehensive methodology that integrates qualitative and quantitative comparative analyses of normal variants and physiological artifacts in simultaneously recorded MEG and EEG data. With this, we aim to enhance the understanding of the unique and complementary characteristics of the two time-resolved noninvasive functional brain imaging modalities, offering insights into their optimal and informed use in clinical and research contexts.

## Methods

Magnetoencephalography (MEG) and electroencephalography (EEG) were registered using a 366-channel hardware and software system ”VectorView” (Elekta Neuromag Oy, Finland) (204 gradiometers, 102 magnetometers, 60 EEG channels). The data in both modalities were recorded concurrently at the sampling rate of 1000 Hz with 0.1- 330 Hz bandwidth. The position of the head relative to the MEG sensor array during the experiment was monitored in real time using special head positioning coils. In our EEG setup, the reference electrode was strategically placed either on the hairline or occasionally on the earlobe to optimize signal clarity and reduce the interference. The ground electrode was positioned on the clavicle. The duration of the experimental session was approximately 80 min.

Ten healthy volunteers (7 females, 3 males) aged 19 to 34 years participated in the study. The inclusion criteria required that the participants had no history of epilepsy, had not undergone surgical procedures that altered the bony structure of the skull, and did not have any implanted metal constructs. Pregnancy was also an exclusion criterion. This study was conducted in accordance with the Declaration of Helsinki; the research adhered to the guidelines and regulations of the institutional and ethical review board of the Higher School of Economics. All participants provided a written informed consent and agreed to participate.

All participants experienced sleep deprivation, with their sleep duration reduced by three hours from the typical amount the previous night. During the recording, the volunteers followed the sequence of the following behavioral probes: eyes open (2 min); eyes closed (2 min); open-closed eyes for 5 seconds (1 min); eye movement in the vertical plane (30 sec); eye movement in the horizontal plane (30 sec); swallowing (ingestion)(30 sec); lip licking (30 sec); coughing (30 sec); smiling (30 sec); sliding head movements in the horizontal plane (30 sec); chewing (30 sec); hyperventilation (2 min).

The preliminary data examination included manual identification of common physiological patterns and scheduled artifacts from EEG data performed by a single clinical neurophysiologist (A.Sh.), then selection of MEG epochs corresponding to these patterns. The duration of epochs was chosen according to the length of the pattern. We focused on several key morphophysiological patterns including alpha spindles, sensorimotor rhythm, PSWoY, vertex patterns, sleep spindles, K-complexes, POSTs, blink artifacts, chewing artifacts, artifacts originating from the forced smile, artifacts originating from the sliding head movements, and ingestion artifacts.

Then, we assessed the spatial and time domain characteristics by aligning and averaging these epochs for each participant. We refrained from implementing complex preprocessing techniques on both EEG and MEG data to directly assess their sensitivity to artifacts and inherent physiological characteristics. The preprocessing was limited to bandpass filtering within the 1-40 Hz range, using an averaged reference for EEG, and interpolating occasionally bad EEG channels, all without specifically targeting artifacts caused by physiological factors. Additionally, we applied the MaxFilter procedure to MEG data: this approach separates the magnetic fields produced by sources within the brain from those produced by the external noise sources. This allows for data standardization across sessions and subjects.

Our quantitative analysis included the averaging procedure of the labeled patterns of stereotypic morphology or the time-frequency decomposition. Time-frequency responses (TFRs) were analyzed for those patterns that were characterized by oscillatory activity (alpha rhythmic activity, SMR, sleep spindles). For this purpose, we utilized complex Morlet wavelets. We analyzed the frequency range from 3 Hz to 30 Hz, since this range covered the dominant frequency components present in the given patterns. The number of cycles in the wavelets was set to be half of their respective central frequency, allowing for an adaptive resolution that increases with the wavelet’s central frequency. Additionally, we applied z-score standartization with respect to the baseline pre-event interval.

Furthermore, we source localized the physiological patterns using the minimum norm estimate (MNE) approach (Hämäläinen and Ilmoniemi, 1994) and visualized the results using the standard FreeSurfer MRI template ’fsaverage’. The noise covariance matrix was calculated from the baseline period, consisting of several seconds preceding the onset of each pattern. The regularization parameter was determined using the L-curve method (Hansen, 1999; Cultrera and Callegaro, 2020) to balance data fidelity and the norm of the solution. Exemplar L-curves corresponding to the chosen morphophysiological patterns are prersented in Fig. 1). For the oscillatory patterns identified by means of the time-frequency decomposition, we applied the inverse operator to the decomposed complex-valued data from the specific time-frequency window and visualized the absolute value of the result.

**Figure 1:**
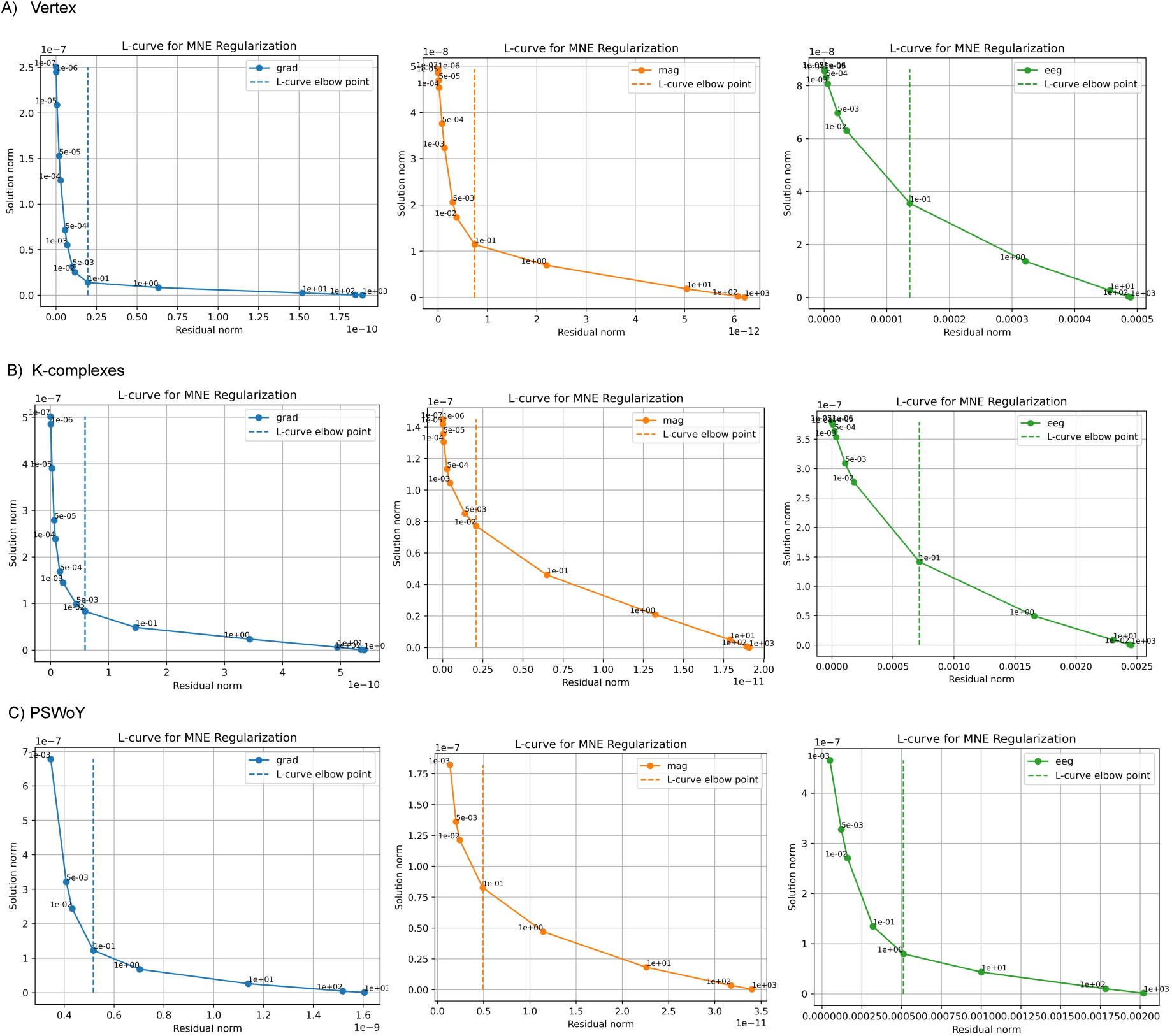
L-curve plots for determining the regularization parameter in the minimum norm estimate (MNE) source localization. Each curve shows the trade-off between data fidelity and the norm of solution, with the optimal parameter chosen at the point of maximum curvature (elbow of the curve): (A) Vertex, (B) K-complexes, (C) PSWoY

Of particular interest is comparing an artifact and physiologically normative brain event that appear morphologically similar in the non-invasive EEG and MEG measurements. To this end we used the group independent components analysis (ICA) (Lee and Lee, 1998) and explored the independent components (ICs) resulting from the spatial decomposition of epochs containing the vertex waves, K-complexes, eye-blink and ingestion artifacts in ten participants. All of these events are characterized by the morphologically similar slow-wave amplitude oscillations.

We used the approach described in (Ossadtchi et al., 2013) to identify the ICs that contain the target events. The ICA decomposition was applied to the concatenated epochs within the +- 2 seconds window around the peak of the pattern for each sensor type separately. The signal was filtered in the range of 1-40 Hz before epoch extraction.

The number of independent components was set to 15. Normalized mutual information (MI) between each IC and the reference signal was then calculated.

As the reference signal we used the global field power (GFP) profile, calculated by averaging across all epochs and all EEG electrodes (Fig. 2) convolved with the binary indicator sequence reflecting the onset of the correspoding activity pattern (the peaks of the vertex waves, K-complexes, eye blinks or ingestion artifacts). This GFP served as the reference for identifying relevant components associated with the pattern. We used normalized mutual information to determine which components contained activity pertaining to the GFP reference and which not, following the approach outlined in (Ossadtchi et al., 2013). To further validate these findings, we conducted a statistical test where we randomized the pattern onset moments and performed 100 repetitions. The statistical significance of each component’s match to the reference was assessed using a conservative threshold of 0.01 (instead of the standard 0.05) to minimize the risk of false positives, given the high amplitude of the patterns under study.

**Figure 2:**
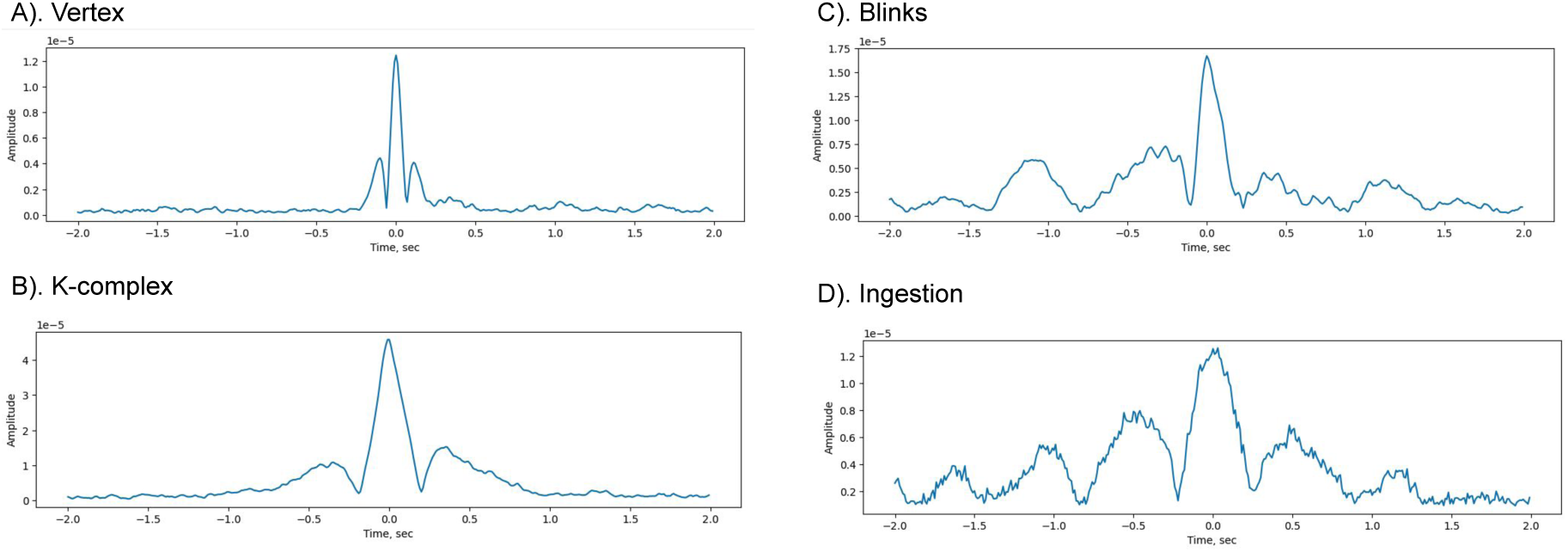
Reference patterns for computing mutual information: global field power of the (A) vertex waves, (B) K-complexes, (C) eye blinks, and (D) ingestion

We conducted a detailed signal-to-noise ratio (SNR) analysis for the identified morphophysiological patterns using a methodology that accounts for overlapping sensor lead fields, as described in (Iivanainen et al., 2021; Petrenko et al., 2021). This approach incorporates the concept of total information capacity to quantify the sensitivity of each sensor type to neuronal signals while mitigating the influence of noise. The mathematical framework for SNR estimation is detailed in the Appendix, where the formulation considers the covariance structure of the sensor measurements and source activity.

In what follows we present the results of applying this methodology to the simultaneously recorded EEG and MEG data in 10 healthy subjects. To enhance the comparative analysis, we have devised a multifaceted data presentation strategy that integrates examples of simultaneous MEG/EEG recordings with quantitative processing and source localization.

## Results

### Physiological patterns

During physiological wakefulness the alpha spindles were induced by closing the participant’s eyes (Fig. 3). As expected this morphophysiological pattern is characterized by occipital topography. MEG and EEG time series (Panel A) captured these spindles with varying degrees of spatial precision. MEG recordings revealed more focal and spatially diverse patterns compared to the broader distributions observed in EEG. Interestingly, while MEG magnitometers show identical to EEG lateralization of the alpha rhythm source to the left hemisphere, the gradiometers highlight bilateral structure of the cortical alpha rhythm sources.

**Figure 3:**
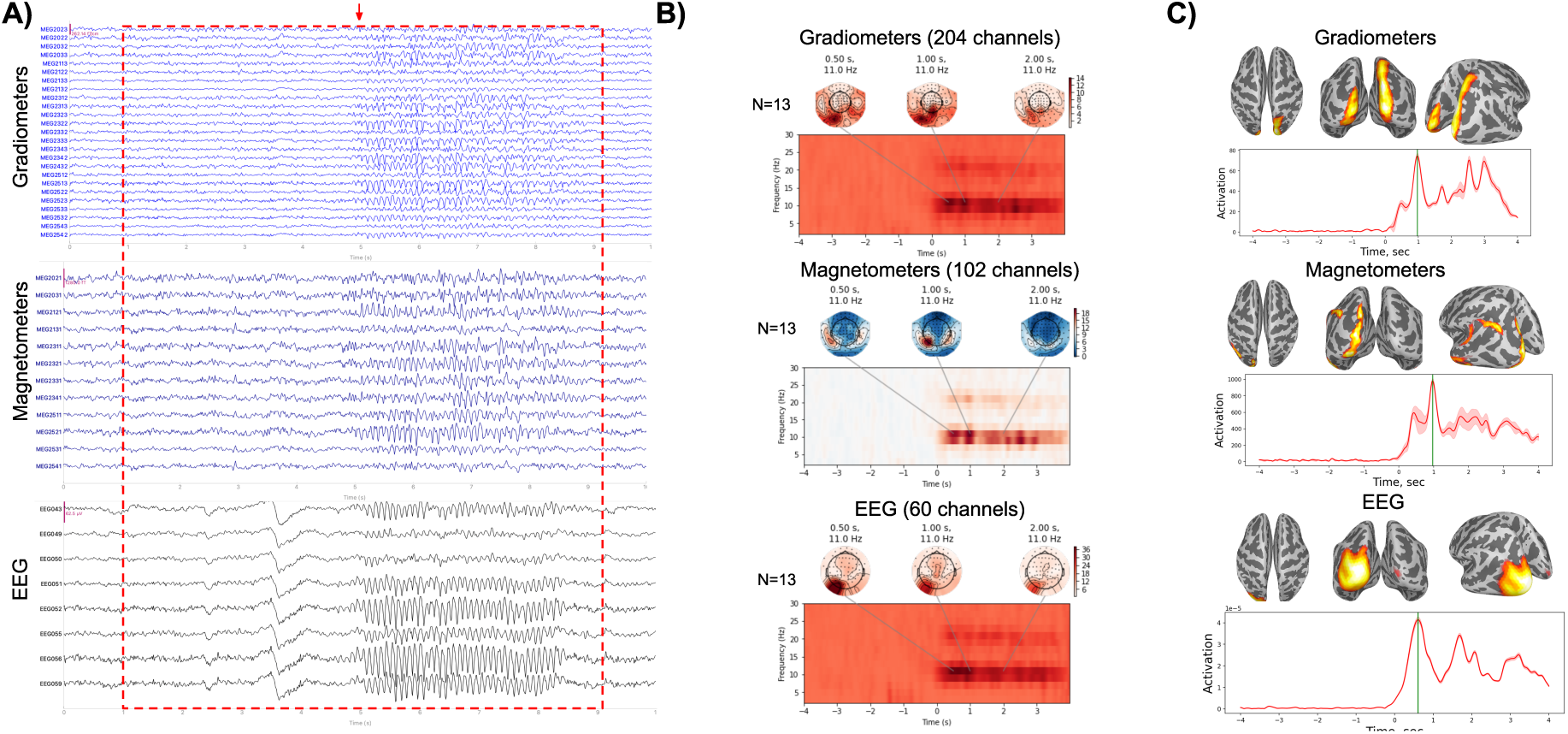
Time, time-frequency domain and spatial characteristics of alpha spindles. (A) Example of patterns in the multi-channel MEG/EEG time series from the right occipital regions. (B) The TFR plots representing the pattern averaged over the subjects (13 epochs), accompanied by topographies showcasing the pattern’s most pronounced expression. The red rectangle in the plot (A) depicts the epoch with the time window equal to the one demonstrated in (B). The red arrow marks the onset of the pattern. (C) Cortical source distributions corresponding to teh spectral components centered around f = 11 Hz.

Sensorimotor rhythm (SMR) (9-11 Hz) was observed in fronto-central EEG sites demonstrating the clear lateralization in the right sensorimotor regions in all three sensor types (Fig. 4). In the time series, all three modalities—gradiometers, magnetometers, and EEG—displayed pronounced arc-like features. These arcs arise from the presence of harmonics, which are integer multiples of the fundamental frequency of the SMR. While EEG indicated a broader distribution biased towards the temporo-parietal junction, MEG—particularly the magnetometers—revealed more localized patterns centered over the sensorimotor regions on both side of the central sulcus. We can also see that localizations delivered by the source analysis of EEG appears to be more central and biased with respect to the sources obtained from the MEG data which may be explained by the inherently lower accuracy of the EEG forward model as compared to that of MEG.

**Figure 4:**
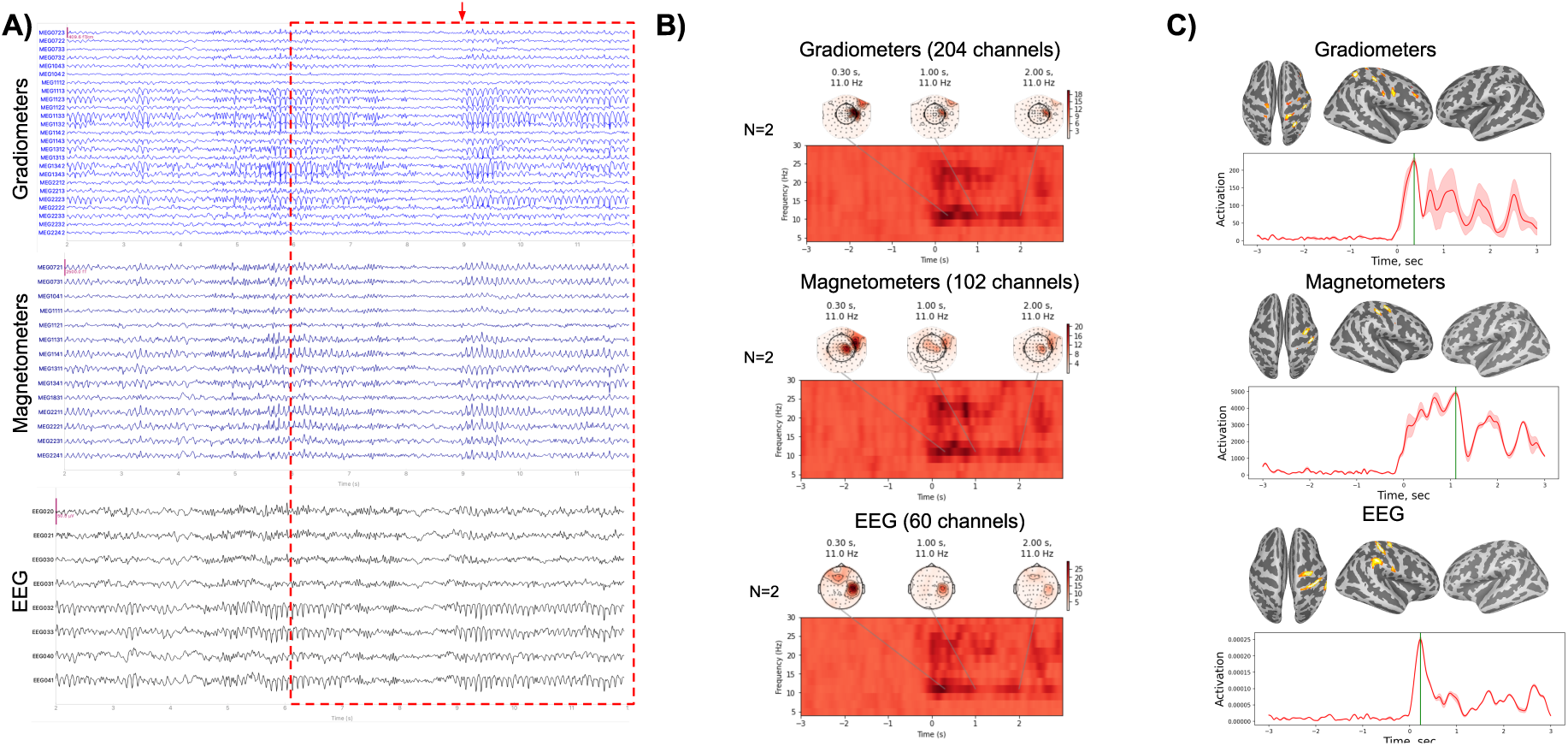
Dynamical and spatial characteristics of the SMR. (A) Example of the patterns in multi-channel MEG/EEG time series from the right central regions. (B) The TFR plots representing the pattern averaged over the subjects (2 epochs), accompanied by topographies showcasing the pattern’s most pronounced expression. The red rectangles on the plot (A) depicts the epoch with a time window equal to the one demonstrated in (B). The red arrow marks the onset of the pattern. (C) The visualization of source modeling of the TFR component at 11 Hz.

Within the main rhythm’s structure, bi-occipital delta waves were occasionally recorded in one subject, without disrupting the underlying rhythmicity, indicative of the physiological pattern known as ”posterior slow waves of youth” (PSWoY) (Fig. 5). In EEG, PSWoY is characterized by a more pronounced peak amplitude, reflecting the modality’s heightened sensitivity to radially oriented cortical sources. Butterfly plots and topographies (Panel B) further highlighted the spatial distribution of PSWoY, with MEG providing more localized representations. In EEG, PSWoY exhibited a wider spread, consistent with the modality’s lower spatial resolution and the increased sensitivity to radially oriented cortical sources as compared to MEG. For the PSWoY localizations based on both MEG sensor types appear highly concordant and show spatially resolved picture indicating the presence of 2-3 connected cortical regions in each of the hemisphere. Going back to the timeseries (Fig. 5.A) one could also notice that moving from EEG to MEG gradiometers we are observing first global and not very well spatially resolved picture (EEG) followed by a more detailed view on the distributed spatial-temporal dynamics yielded by MEG magnetometers and finally arrive at a very detailed and high-frequency (both spatially and temporally) picture revealed by MEG gradiometers. Noteworthy is that the combination of EEG and MEG allows to easily avoid an erroneous decision of classifying the sharp spike-wave pattern seen in several gradiometer channels as an interictal spike.

**Figure 5:**
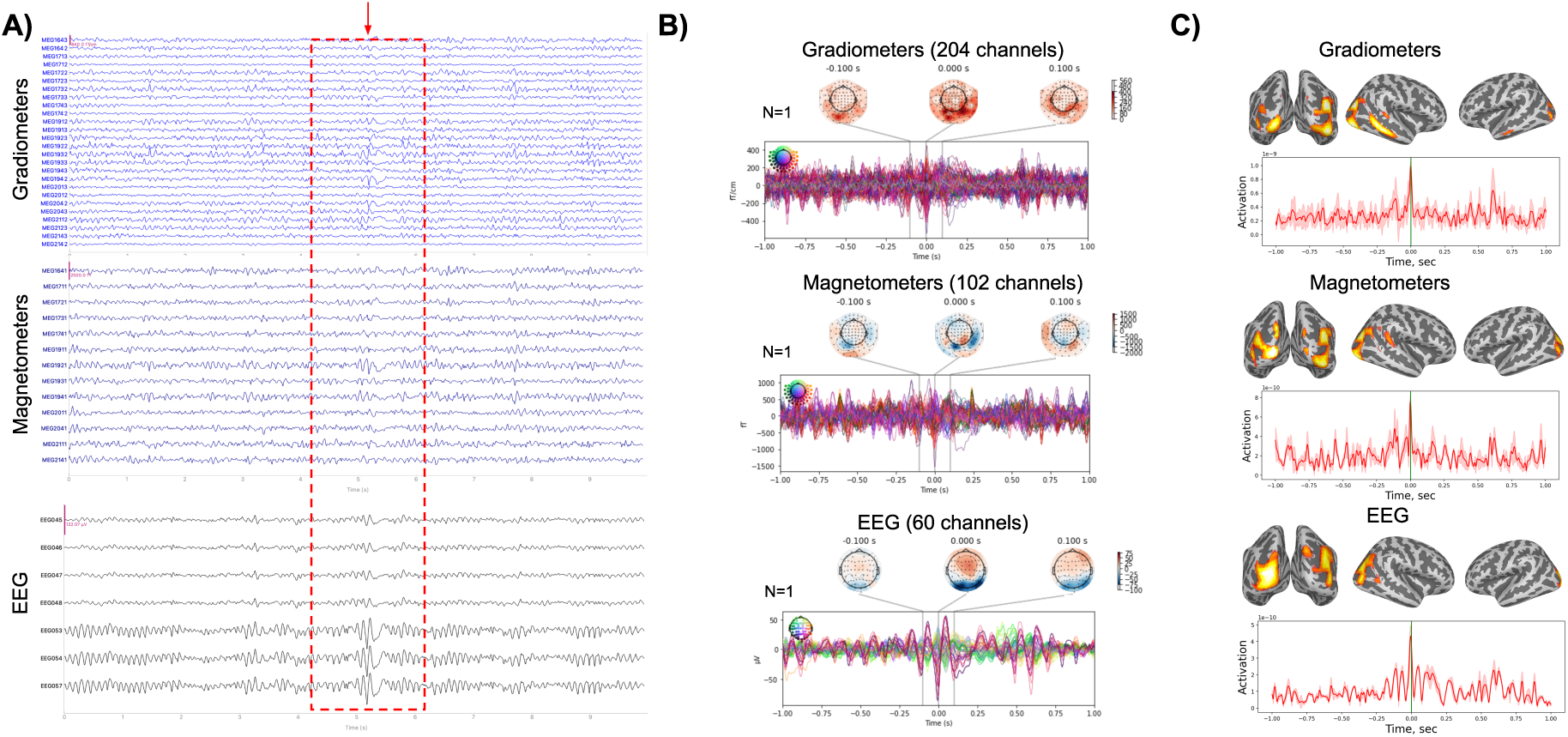
Dynamical and spatial characteristics of PSWoY. (A) Example of the patterns in multi-channel MEG/EEG time series from the left occipital regions. (B) The butterfly plots representing the pattern (1 epoch), accompanied by topographies showcasing the pattern’s most pronounced expression. The red rectangle in the plot (A) depicts the epoch with the time window equal to the one demonstrated in (B). The red arrow marks the peak of the pattern. (C) The visualization of source modeling of the pattern.

All volunteers were instructed to transition into sleep. The first and second stages of non-REM sleep were observed in all participants, marked by a progressive decrease in alpha rhythm, the emergence of low-amplitude, mixed-frequency rhythmicity, the appearance of vertex potentials (Fig. 6), the occurrence of sleep spindles (Fig. 7) in the anterior and vertex sections, the registration of K-complexes (Fig. 8) - high-amplitude biphasic or triphasic potentials predominantly in the anterior sections, and positive occipital sharp transient of sleep (POSTs) (Fig. 9).

**Figure 6:**
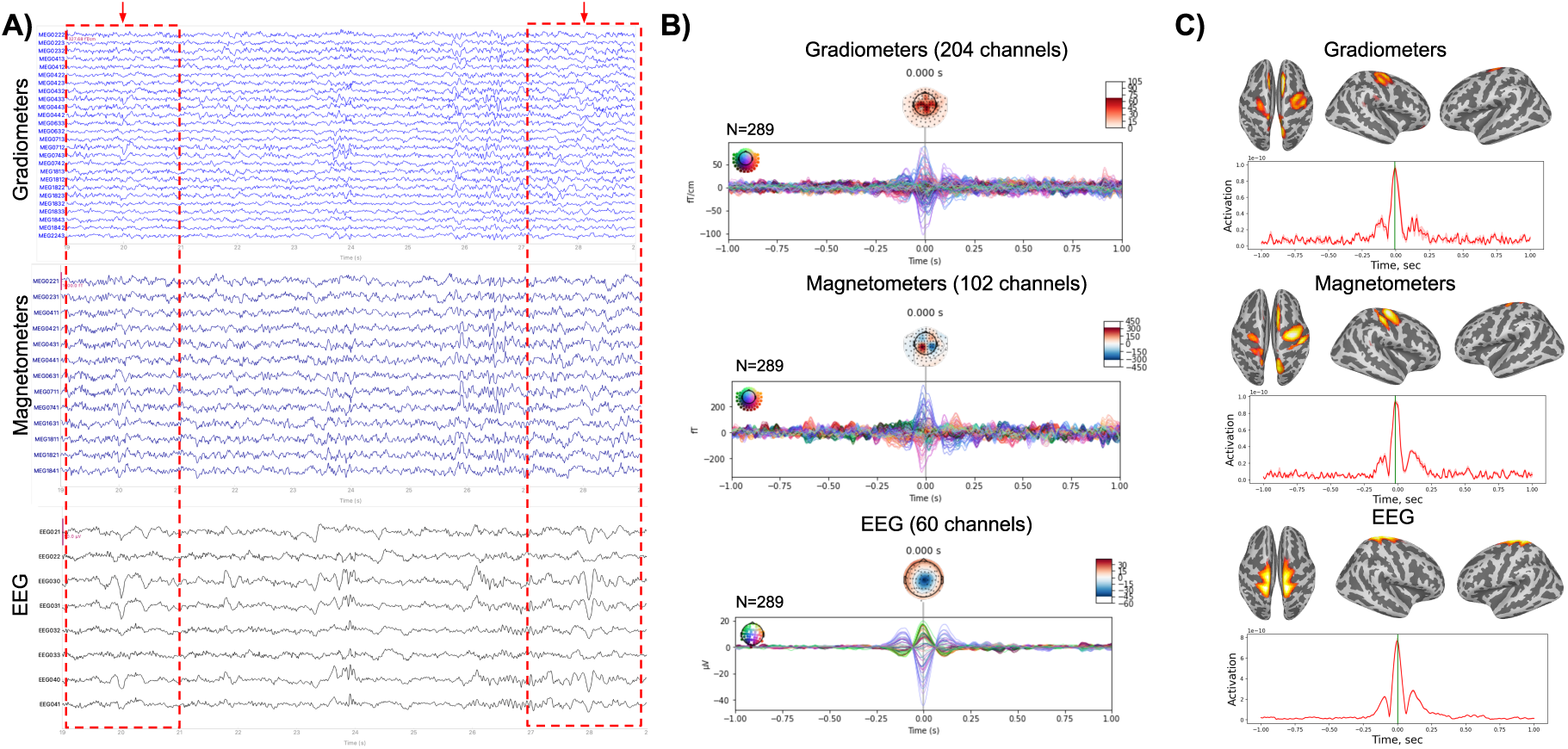
Dynamical and spatial characteristics of vertex patterns. (A) Example of the patterns in multi-channel MEG/EEG time series from the left central regions. (B) The butterfly plots representing the pattern averaged over the subjects (289 epochs), accompanied by topographies showcasing the pattern’s most pronounced expression. The red rectangles in the plot (A) depict 2 epochs with the time window equal to the one demonstrated in (B). The red arrows mark the peak of the pattern. (C) The visualization of source modeling of the averaged pattern.

**Figure 7:**
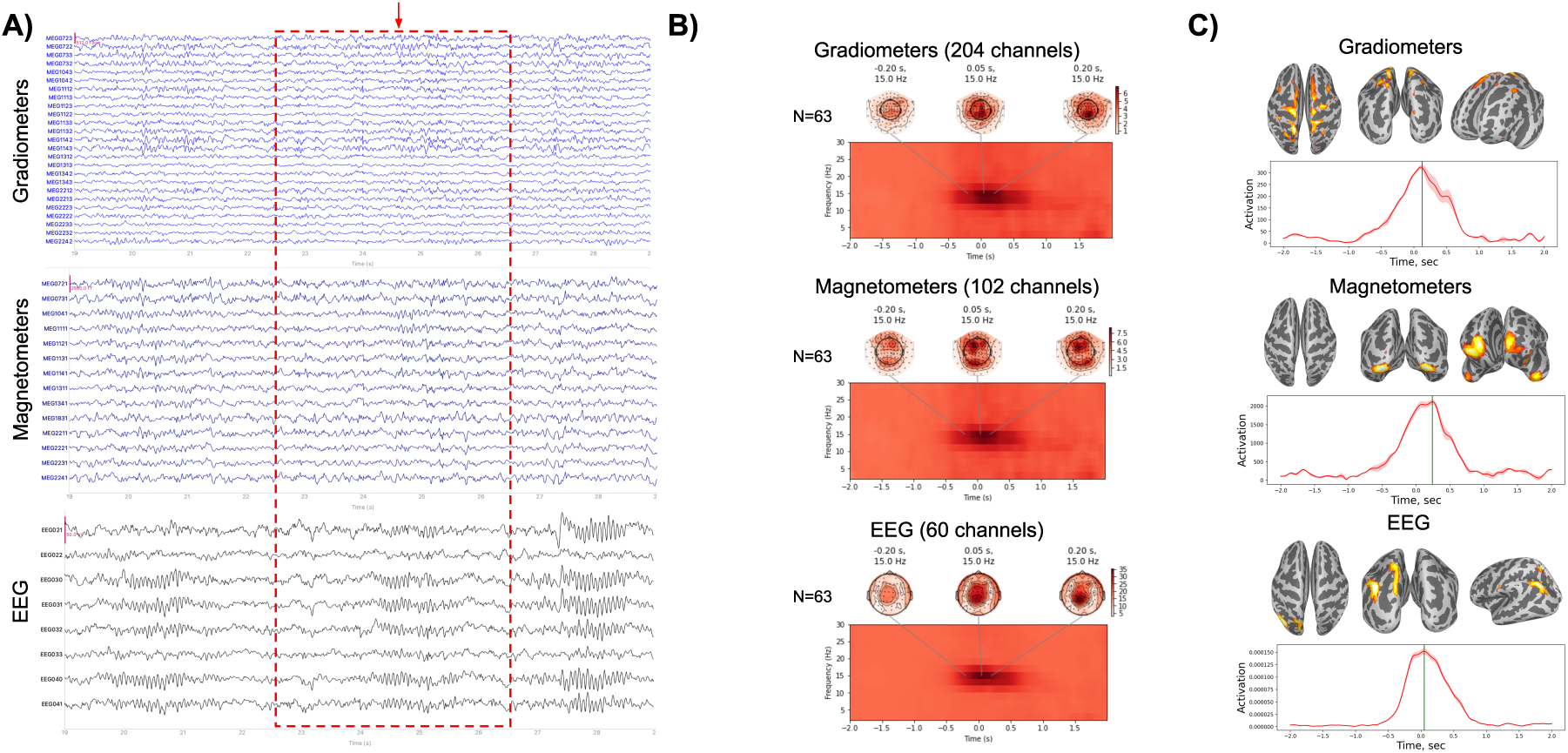
Dynamical and spatial characteristics of sleep spindles. (A) Example of the patterns in multi-channel MEG/EEG time series from the right central regions. (B) The TFR plots representing the pattern averaged over the subjects (63 epochs), accompanied by topographies showcasing the pattern’s most pronounced expression. The red rectangle in the plot (A) depicts the epoch with the time window equal to the one demonstrated in (B). The red arrow marks the peak of the pattern. (C) The visualization of source modeling of the averaged pattern at 15 Hz.

**Figure 8:**
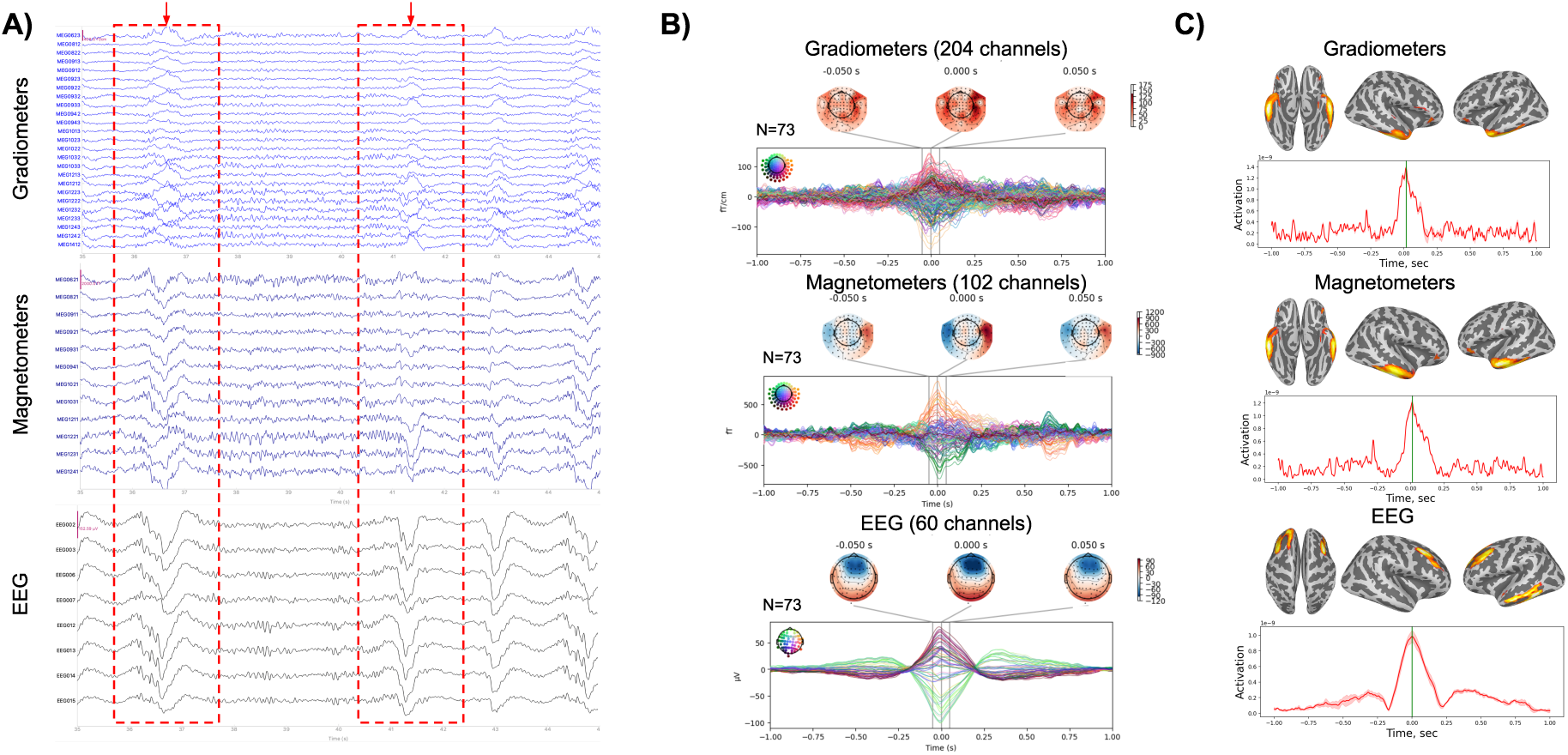
Dynamical and spatial characteristics of K-complexes. (A) Example of the patterns in multi-channel MEG/EEG time series from the right frontal regions. (B) The butterfly plots representing the pattern averaged over the subjects (73 epochs), accompanied by topographies showcasing the pattern’s most pronounced expression. The red rectangles on the plot (A) depict 2 epochs with the time window equal to the one demonstrated in (B). The red arrows mark the peak of the pattern. (C) The visualization of source modeling of the averaged pattern.

**Figure 9:**
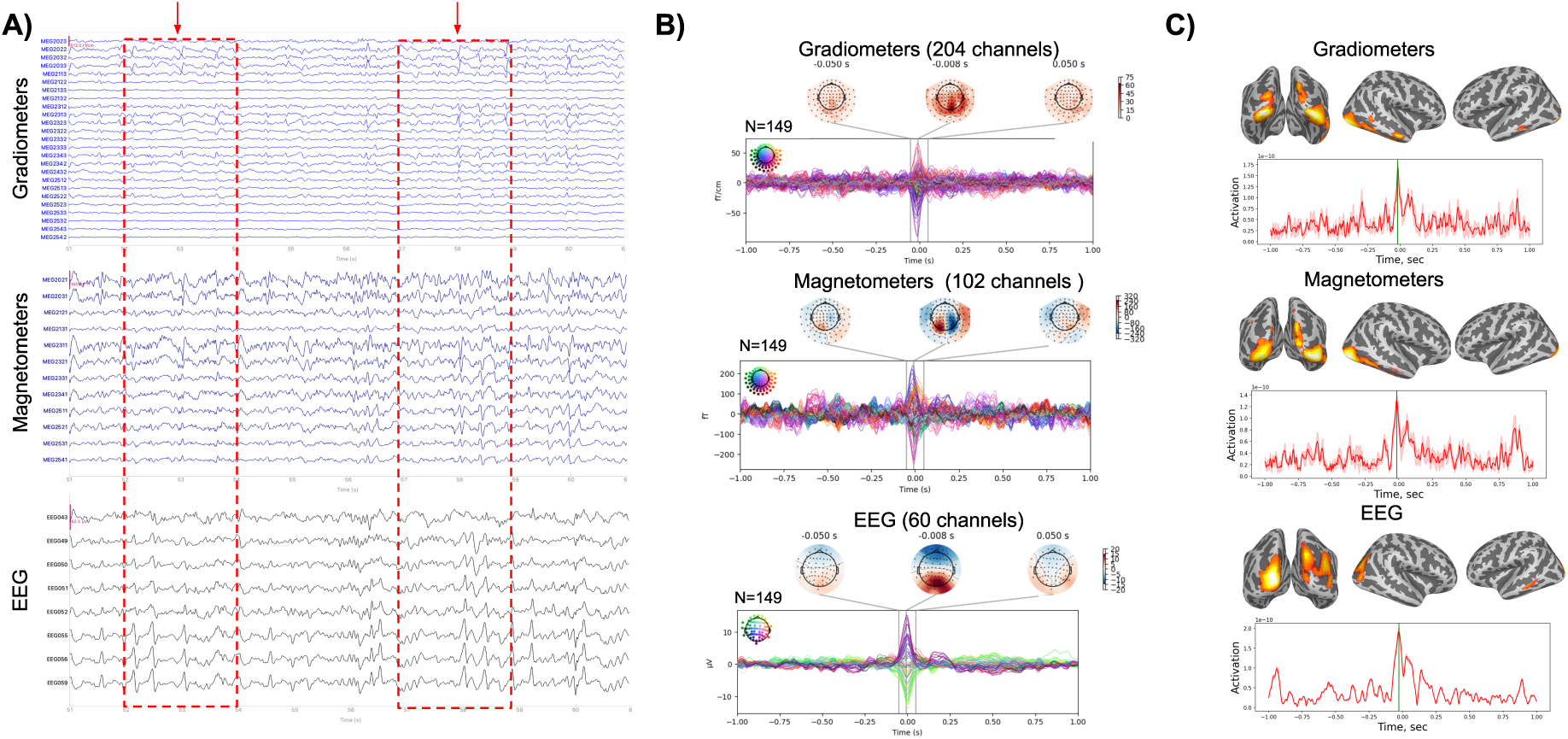
Dynamical and spatial characteristics of POSTs. (A) Example of the patterns in multi-channel MEG/EEG time series from the right occipital regions. (B) The butterfly plots representing the pattern averaged over the subjects, accompanied by topographies showcasing the pattern’s most pronounced expression. The red rectangles in the plot (A) depict the time window equal to the one demonstrated in (B). The red arrows mark the onset of the pattern. (C) The visualization of source modeling of the averaged pattern.

As evident from Fig. 6.B the vertex potentials measured by MEG sensors displayed ’quadrupolar’ topographies, whereas those recorded by EEG channels showed a diffuse spatial pattern corresponding to a radially oriented global source. Source analysis of MEG sleep spindles revealed their fronto-central and parietal origin, while the EEG based sources landed in the superior parietal regions located posterior with respect to the localizations computed form MEG data. Interestingly, the spatial patterns of two framed vertex episodes (Fig. 6.B) appear slightly different. This can be conclude both the analysis of the relative amplitudes of spatial distributions among EEG channels but more vividly using the MEG timeseries. The first episode is seen very well in all three channels types with MEG being more spatially resolved. At the same time the second vertex episode is very weakly captured by the selected MEG channels which may either speak of its neuronal source having dominantly radial orientations or may one more time emphasize a superior spatial resolving power of MEG if this episode is seen well on another subset of channels.

Sleep spindles were predominantly recorded during stage 2 of non-REM sleep and were clearly visible in EEG signals (Fig. 7). The time-frequency representation (Fig. 7, B) revealed a sharp increase in power centered around 15 Hz in all three sensor types. Among them, magnetometers exhibited the most spatially specific pattern, with spindle activity predominantly localized in fronto-parietal regions.

The spatial characteristics of K-complexes in MEG were more complex and localized to the inferior-anterior temporal regions including the temporal poles. In EEG, the K- complexes exhibited a typical central topography but source localization also revealed activation in the lower portions of the temporal lobe, more diffuse and located superior and posterior w.r.t to the source distribution computed from MEG data. In MEG, the K-complexes displayed a more symmetric topography, with gradiometers and magnetometers revealing bilateral activity in temporal regions. This spatial distribution can be attributed to MEG’s prevailing sensitivity to the signals generated by tangentially oriented cortical sources, that are less influenced by the radial components dominating EEG recordings.

POSTs were observed most prominently in the occipital regions (Fig. 9). These waveforms were characterized by sharp, monophasic delta-frequency events with a steep ascending phase and a slower descending return, a pattern consistent with known descriptions of POST morphology. Gradiometers captured brief, high-amplitude transients with a clear polarity reversal across nearby channels, suggesting a localized tangential dipole generator. EEG showed the POSTs as sharp waves with a wider spatial spread.

### Artifacts

During physiological wakefulness recordings the blink artifacts were registered with typical dipolar characteristics in the frontal lobes (see Fig. 10). Noteworthy is that the morphological patterns observed in MEG and EEG (Panel A) appear to be very different for this type of events. For the eye-blinks, unlike for the majority of other patterns presented, the gradiometer MEG channel timeseries appear more similar than their magnetometer counterparts. In the gradiometer channels, unlike it is the case with magnetometers, we can observe sharp step-like patterns similar to those dominating the EEG recordings.

**Figure 10:**
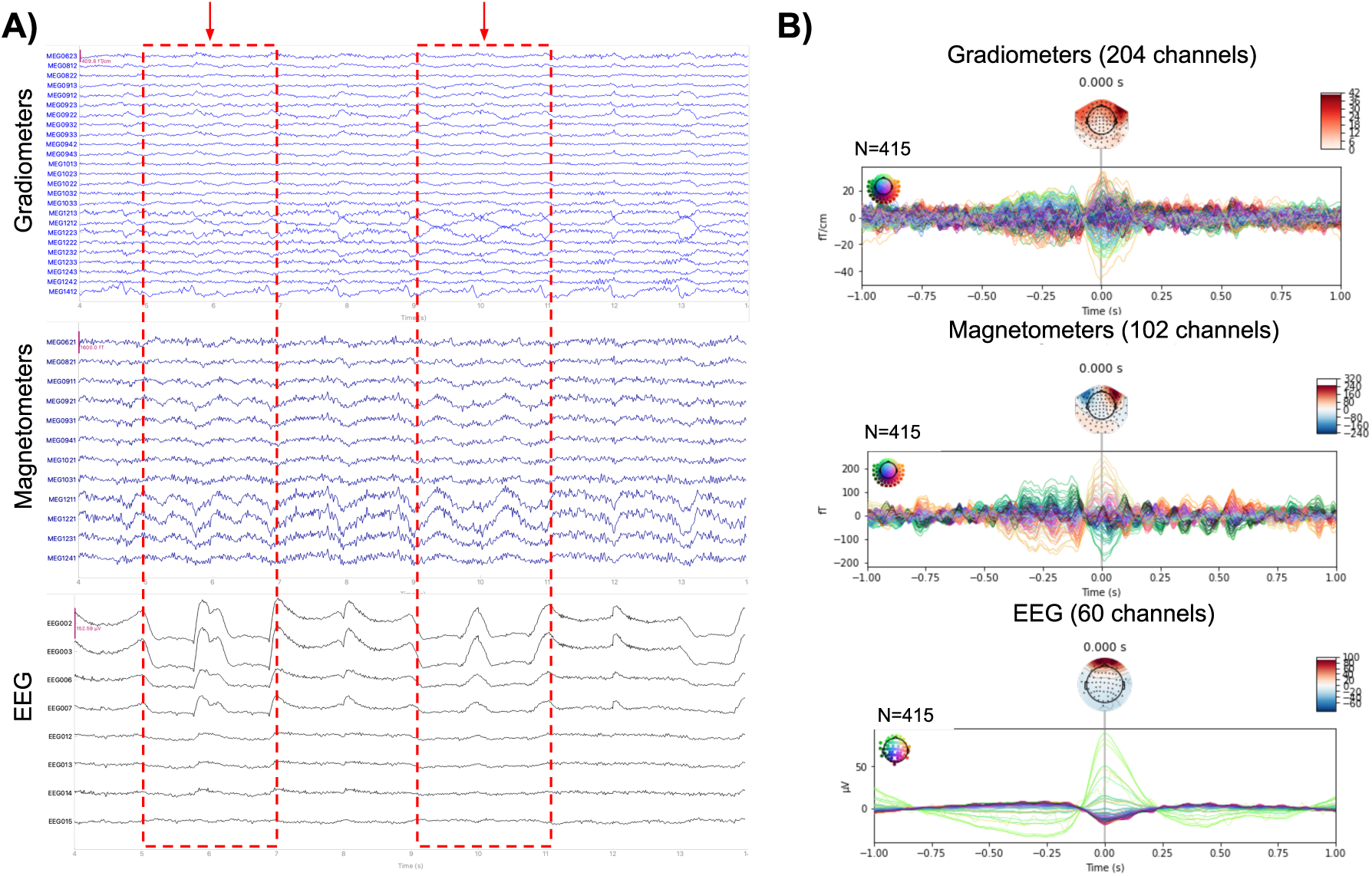
Dynamical and spatial characteristics of blink artifacts. (A) An example of the patterns in multi-channel MEG/EEG time series from the right frontal regions. (B) The butterfly plots representing the pattern averaged over the subjects (415 epochs), accompanied by topographies showcasing the pattern’s most pronounced expression. The red rectangles on the plot (A) depict 2 epochs with time windows equal to the one demonstrated in (B). The red arrows marks the peak of the pattern.

During the recording of chewing artifacts (Fig. 11) and forced smile (grin) (Fig. 12), the EEG channels registered pronounced myographic artifacts that dominated the recordings masking the rest of activity. While the gradiometers and EEG were dominated by local muscular activity, the magnetometers reflected a large dipolar pattern most likely pertaining to the tongue, that is known to be a dipole oriented in the posterior-anterior direction (Loose et al., 2001). See however Bayram et al. (2022) for additional insights into tongue’s electrical properties.

**Figure 11:**
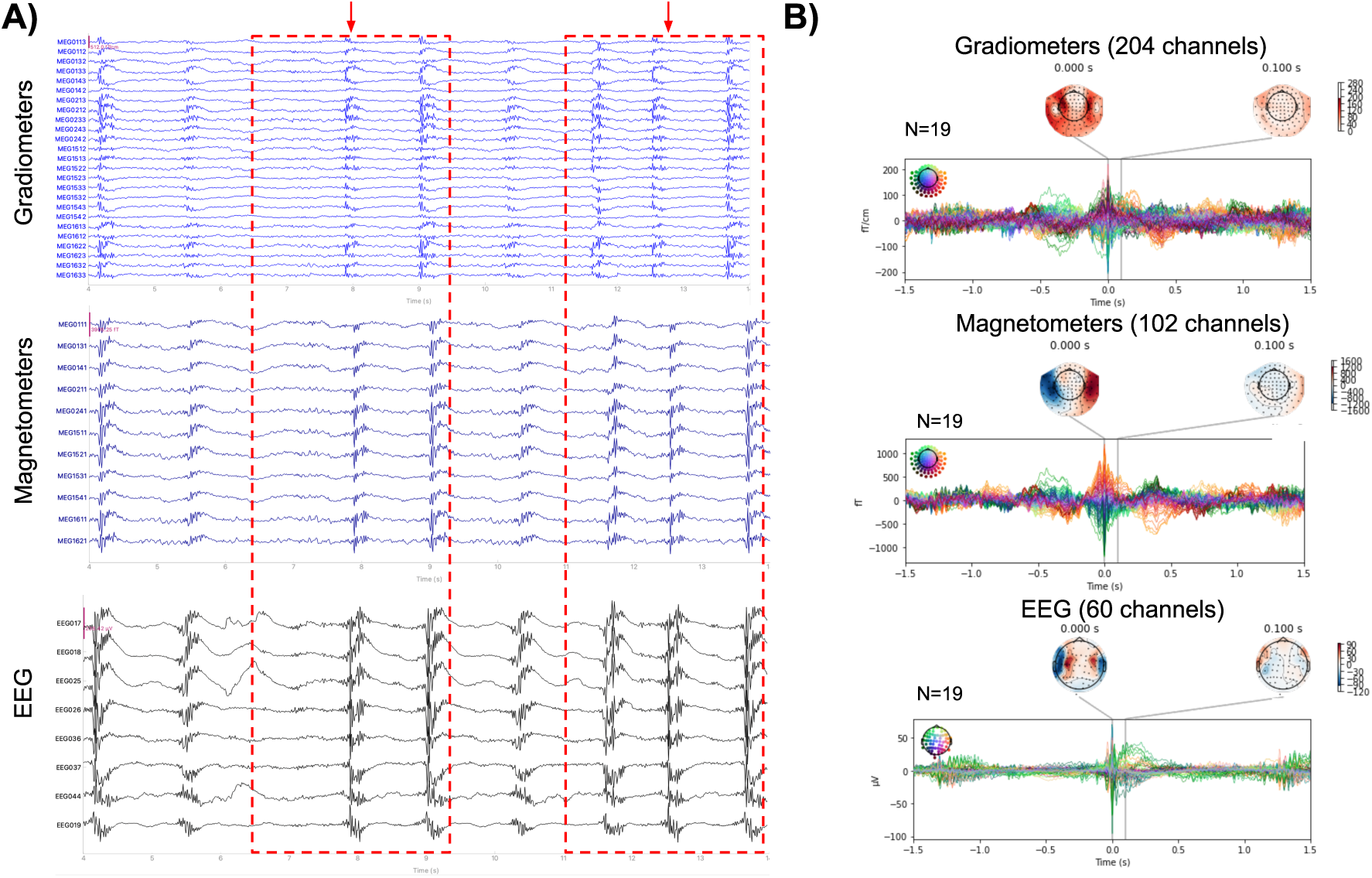
Dynamical and spatial characteristics of chewing artifacts. (A) Example of the patterns in multi-channel MEG/EEG time series from the left temporal regions. (B) The butterfly plots representing the pattern averaged over the subjects (19 epochs), accompanied by topographies showcasing the pattern’s most pronounced expression. The red rectangles in the plot (A) depict 2 epochs with time windows equal to the one demonstrated in (B). The red arrows mark the onset of the pattern.

**Figure 12:**
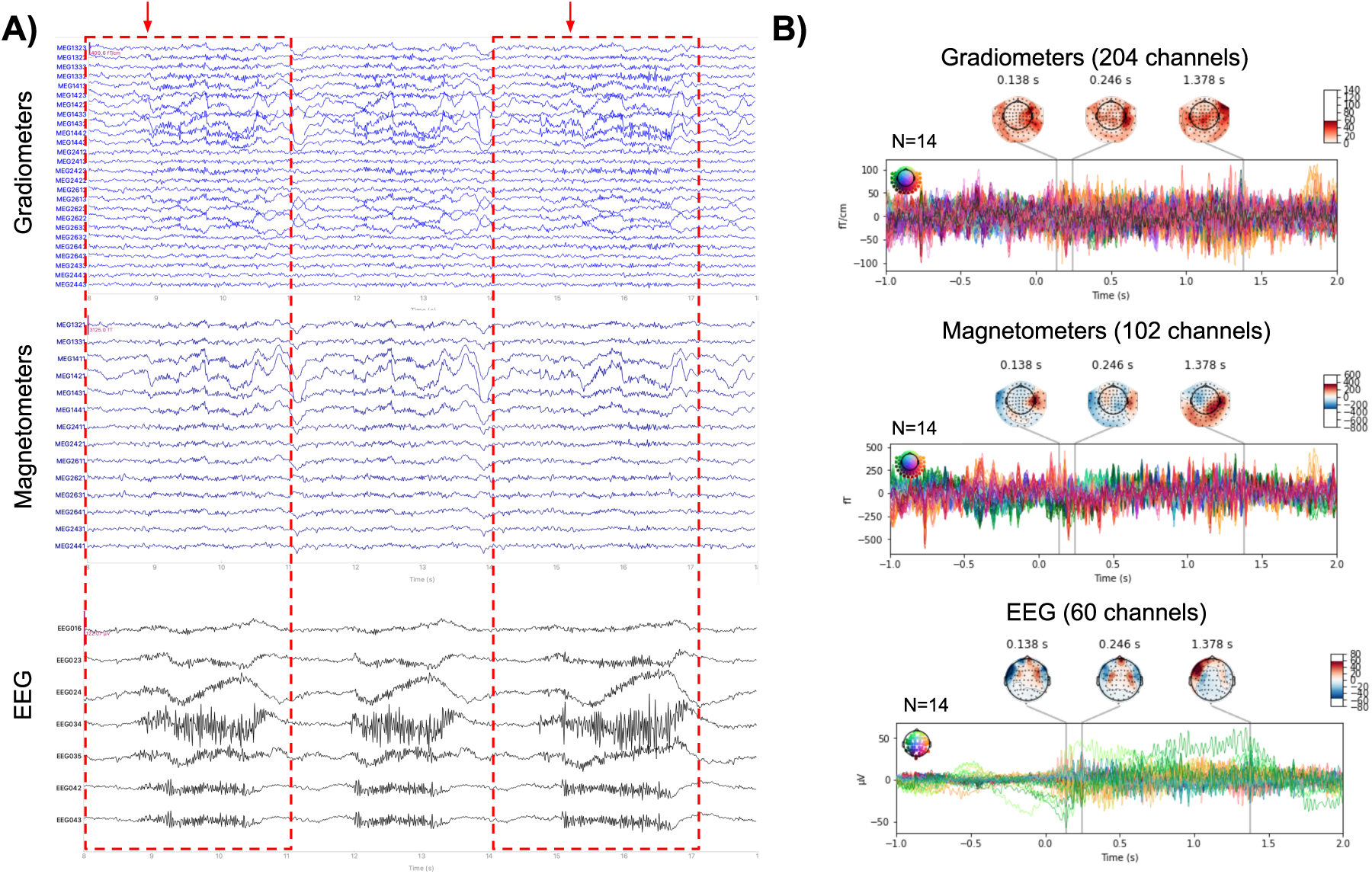
Dynamical and spatial characteristics of artifacts originating from the forced smile. (A) Example of the patterns in multi-channel MEG/EEG time series from the right temporal regions. (B) The butterfly plots representing the pattern averaged over the subjects (14 epochs), accompanied by topographies showcasing the pattern’s most pronounced expression. The red rectangles in the plot (A) depict 2 epochs with the time window equal to the one demonstrated in (B). The red arrows mark the onset of the pattern.

Additionally, we observed the complex artifacts originating from the sliding head movements. They mainly affected frontal and temporal regions and consisted of contributions from high and low frequencies (Fig. 13).

**Figure 13:**
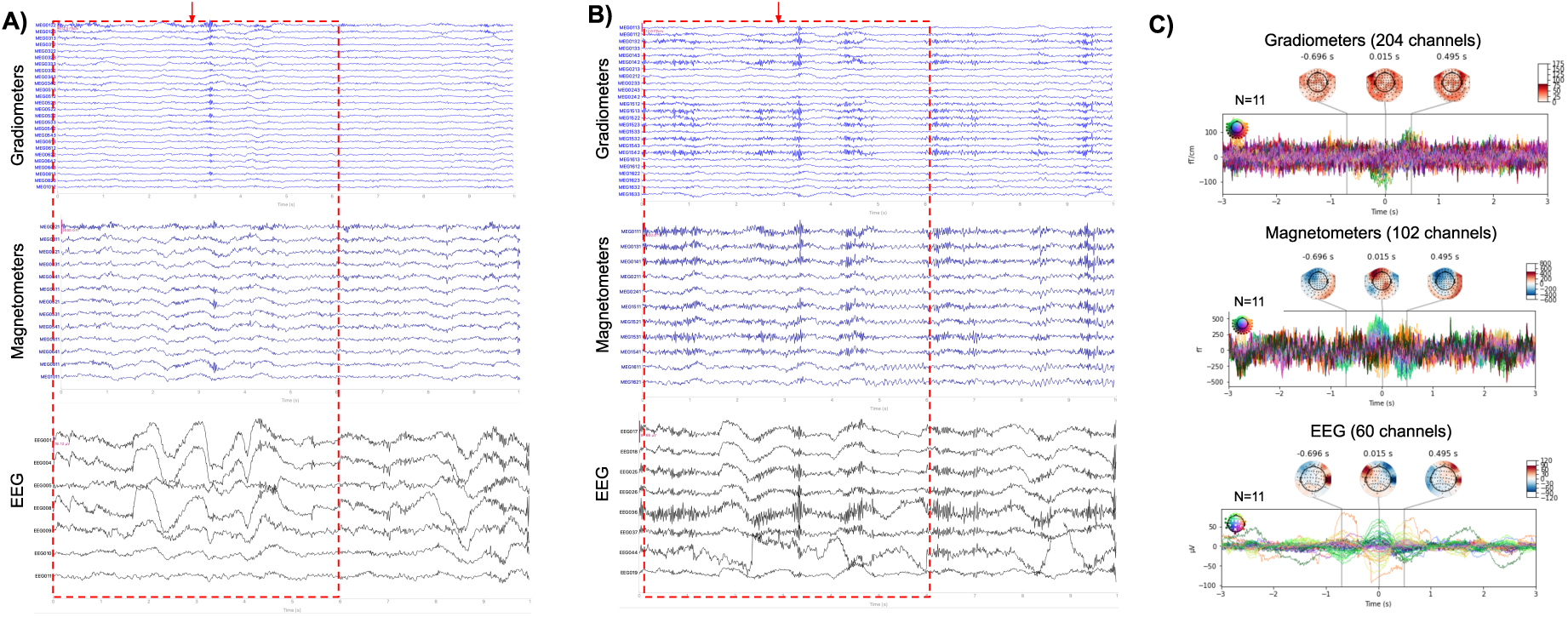
Dynamical and spatial characteristics of artifacts originating from the head movement. Examples of the patterns in multi-channel MEG/EEG time series from the left frontal (A) and left temporal (B) regions. (C) The butterfly plots representing the pattern averaged over the subjects (11 epochs), accompanied by topographies showcasing the pattern’s most pronounced expression. The red rectangles in the plots (A) and (B) depict the epoch with the time window equal to the one demonstrated in (C). The red arrow marks the onset in the time scale of (C).

Finally, the swallowing resulted in high-amplitude rhythmic artifacts affecting the frontal regions (Fig. 14) and having meander-like temporal profiles registered by the sensors of all three modalities.

**Figure 14:**
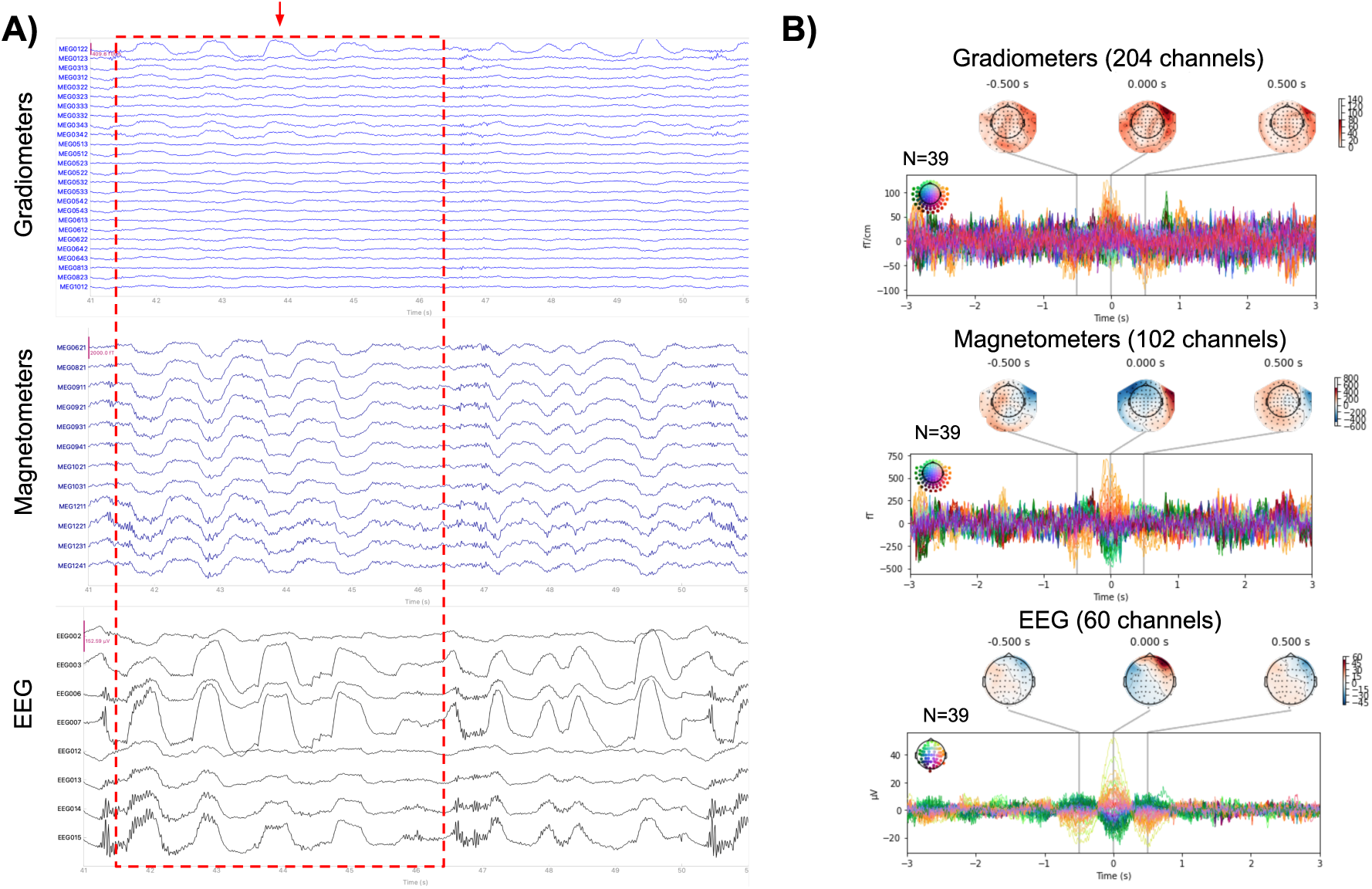
Dynamical and spatial characteristics of artifacts resulting from the swallowing. (A) Example of the patterns in multi-channel MEG/EEG time series from the right frontal regions. (B) The butterfly plots representing the pattern averaged over the subjects (39 epochs), accompanied by topographies showcasing the pattern’s most pronounced expression. The red rectangle in the plot (A) depicts the epoch with the time window equal to the one demonstrated in (B). The red arrow marks the peak of the pattern.

### Quantitative analysis of mutual information

To quantitatively assess the representation of physiological and artifactual patterns in MEG and EEG, we applied ICA to the data across all participants. This approach allowed us to identify statistically significant independent components (ICs) associated with each target pattern— vertex waves, K-complexes, eye blinks, and ingestion artifacts.

For each sensor type (magnetometers, gradiometers, EEG), we computed the mutual information (MI) spectrum between the extracted ICs and the reference signal, which was derived from the global field power (GFP) of the respective pattern (see Section 2). The components whose MI value significantly exceeded the null distribution were considered related to the particular normal variant or artifact .

The MI spectrum for vertex waves (Figure 15) exhibits modality-dependent differences. In EEG (Fig. 15, 3.A), the spectrum is steep, with only one significant component ((Fig. 15, 3.B)), which has the expected central topography. In contrast, in magnetometers (Fig. 15, 1.A), the MI spectrum is more gradual, with four significant components (Fig. 15, 1.B), each corresponding to distinct spatially distributed sources. Some components remain in the vertex region but are oriented orthogonally to each other, while others extend bilaterally towards the temporal regions, shift parietally, or are positioned centrally but closer to the frontal areas. This suggests that MEG captures a more complex underlying structure of the vertex wave, potentially reflecting multiple contributing sources. Additionally, the temporal activation profiles of magnetometer components exhibit substantial variability, particularly in component 4, whose activation is characterized by two peaks. In gradiometers (Fig. 15, 2.A), the MI spectrum is steeper than in magnetometers, with only two significant components ((Fig. 15, 2.B)), yet variability in the spatial distribution of sources is still present. The greater selectivity of gradiometers, compared to magnetometers, may be explained by their higher sensitivity to superficial, tangentially oriented sources, while magnetometers capture a broader field, including deeper sources.

**Figure 15:**
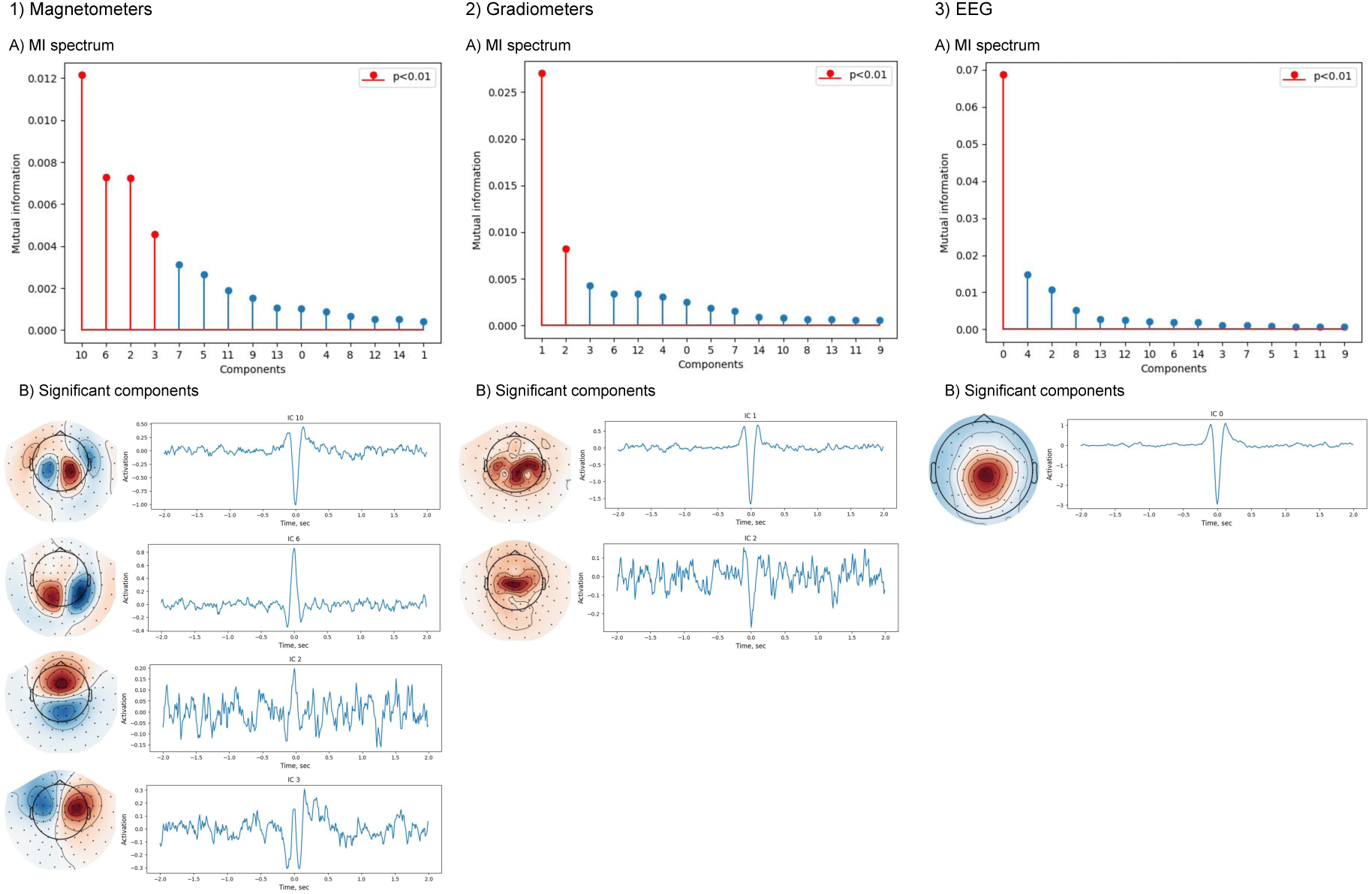
Results of ICA for vertex waves obtained for magnetometers (1), gradiometers (2), and EEG (3): A) MI spectrum illustrating the distribution of mutual information values across ICs. B) Topographies and activation time courses of significant independent components, showing spatial and temporal characteristics of the extracted sources.

It is instructive to compare the topography of the first IC derived from the magnetometer data (Fig. 15, 1.B) to that derived from the EEG signals and depicted in Fig. 15, 3.B. Magnetometer data topography shows two well pronounced bilateral temporal-parietal sources while the topography of the only EEG IC has a structure similar to that observed in the auditory evoked response when two bilaterally symmetric dipoles located in the superior temporal lobes get activated. This may imply that the only EEG’s IC directly corresponds to the first IC derived from the magnetometer data. The rest of the MEG derived components exhibiting significant mutual information with vertex events represent additional information about the spatial-temporal cortical dynamics underlying the vertex waves captured by MEG.

A similar tendency is observed for K-complexes (Figure 16). In EEG (Fig. 16, 3.A), the MI spectrum is steep, with only two significant components (Fig. 16, 3.B), both exhibiting frontal-central topography, consistent with the known localization of K-complexes. In contrast, magnetometers and gradiometers (Fig. 16, 1.A-2.A) reveal a more gradual MI spectrum, with six and five significant components, respectively (Fig. 16, 1.B-2.B). As with vertex waves, these components demonstrate a wider spatial distribution, including frontal, central, and even parietal areas, and exhibit differences in temporal activation profiles. This suggests that while EEG represents K-complexes as a more unitary event, MEG reveals a richer structure, possibly capturing different cortical contributions to this waveform.

**Figure 16:**
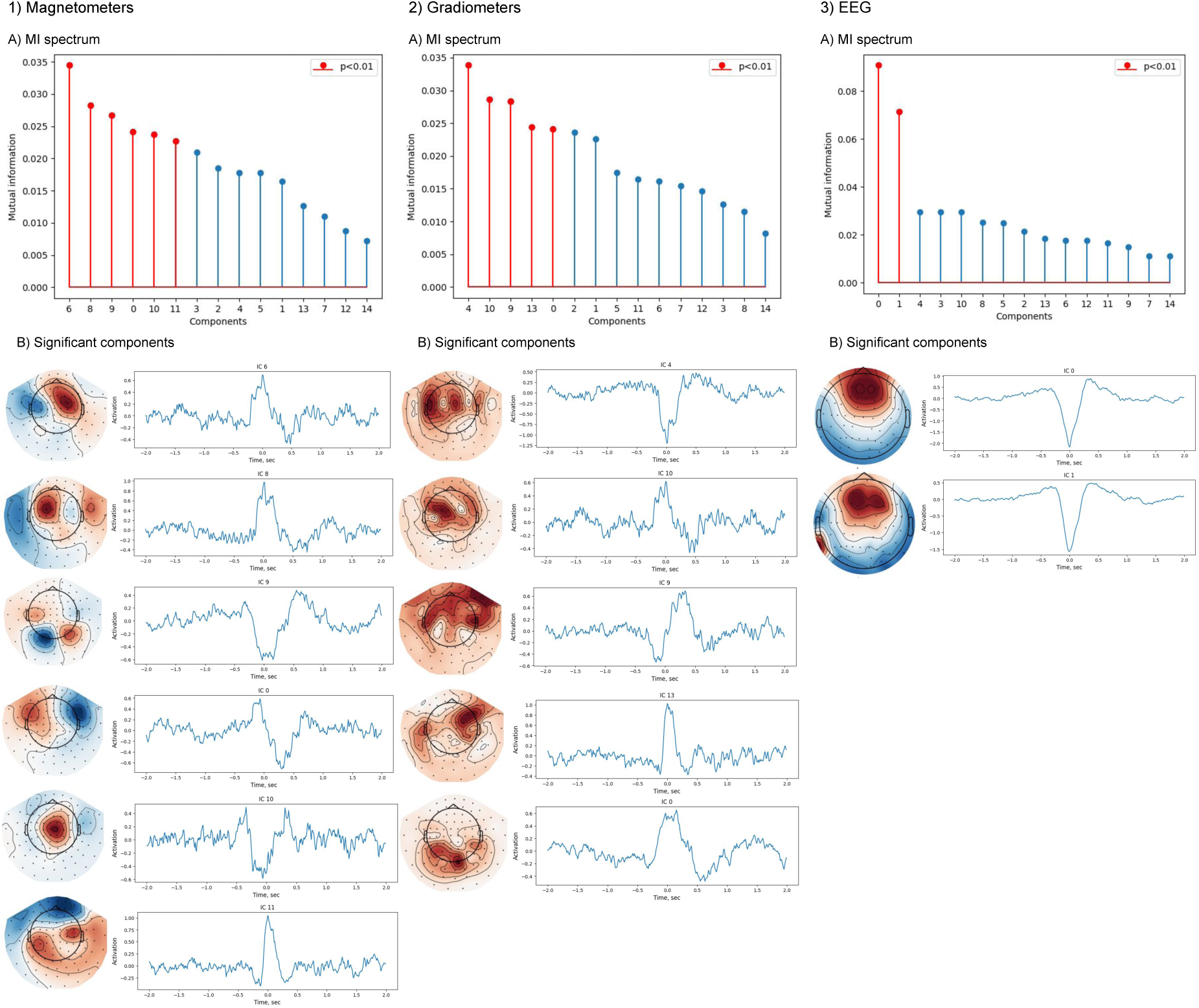
Results of ICA for K-complexes obtained for magnetometers (1), gradiometers (2), and EEG (3): A) MI spectrum illustrating the distribution of mutual information values across ICs. B) Topographies and activation time courses of significant independent components, showing spatial and temporal characteristics of the extracted sources.

For artifacts, the results differ. In the case of eye blinks (Fig. 17), EEG detects multiple significant components (Fig. 17, 3.A-B), all with frontal topography, but with distinct temporal dynamics. In magnetometers and gradiometers (Fig. 17, 1.A-2.A), the MI spectrum is much steeper, with only two significant components in each case (Fig. 17, 1.B-2.B). In magnetometers, one component exhibits the expected frontal topography, while the other appears to originate from a deeper source. gradiometers largely repeat both spatial and temporal patterns observed in magnetometers. Overall, MEG captures information about the artifacts in a set of fewer components than EEG which allows for a more efficient spatial filtering.

**Figure 17:**
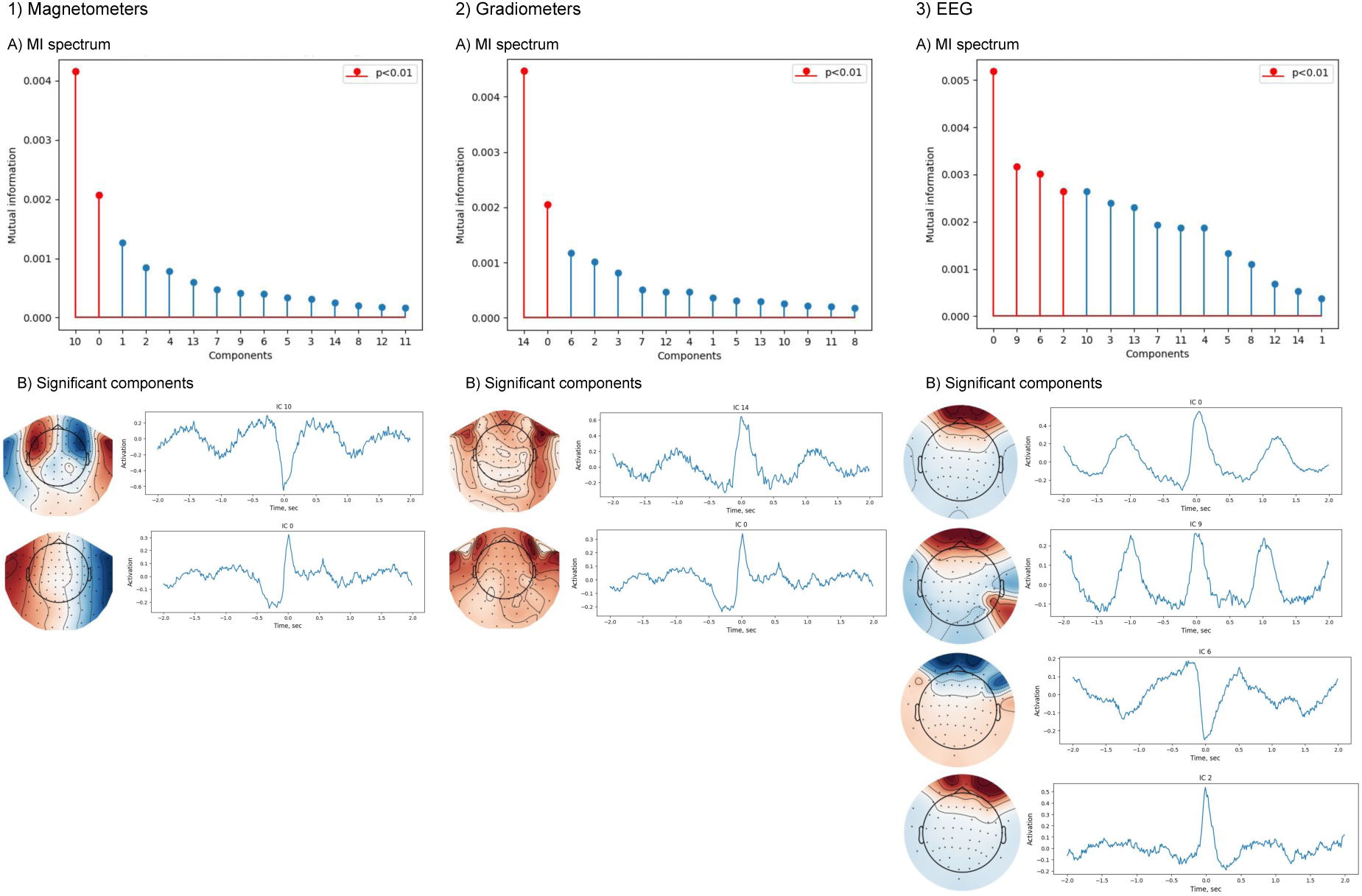
Results of ICA for eye-blink artifacts obtained for magnetometers (1), gradiometers (2), and EEG (3): A) MI spectrum illustrating the distribution of mutual information values across ICs. B) Topographies and activation time courses of significant independent components, showing spatial and temporal characteristics of the extracted sources.

A similar pattern is observed for ingestion artifacts (Figure 18). In EEG (Fig. 18, 3.A-B), three significant components are detected, with sources distributed across the frontal and temporal regions, consistent with the widespread muscle activity associated with swallowing. In magnetometers and gradiometers (Fig. 18, 1.A-2.A), only two significant components are found (Fig. 18, 1.B-2.B), both with frontal topographies. The MI spectrum in MEG is steeper than in EEG, again suggesting that MEG captures a more selective subset of the artifact-related activity.

**Figure 18:**
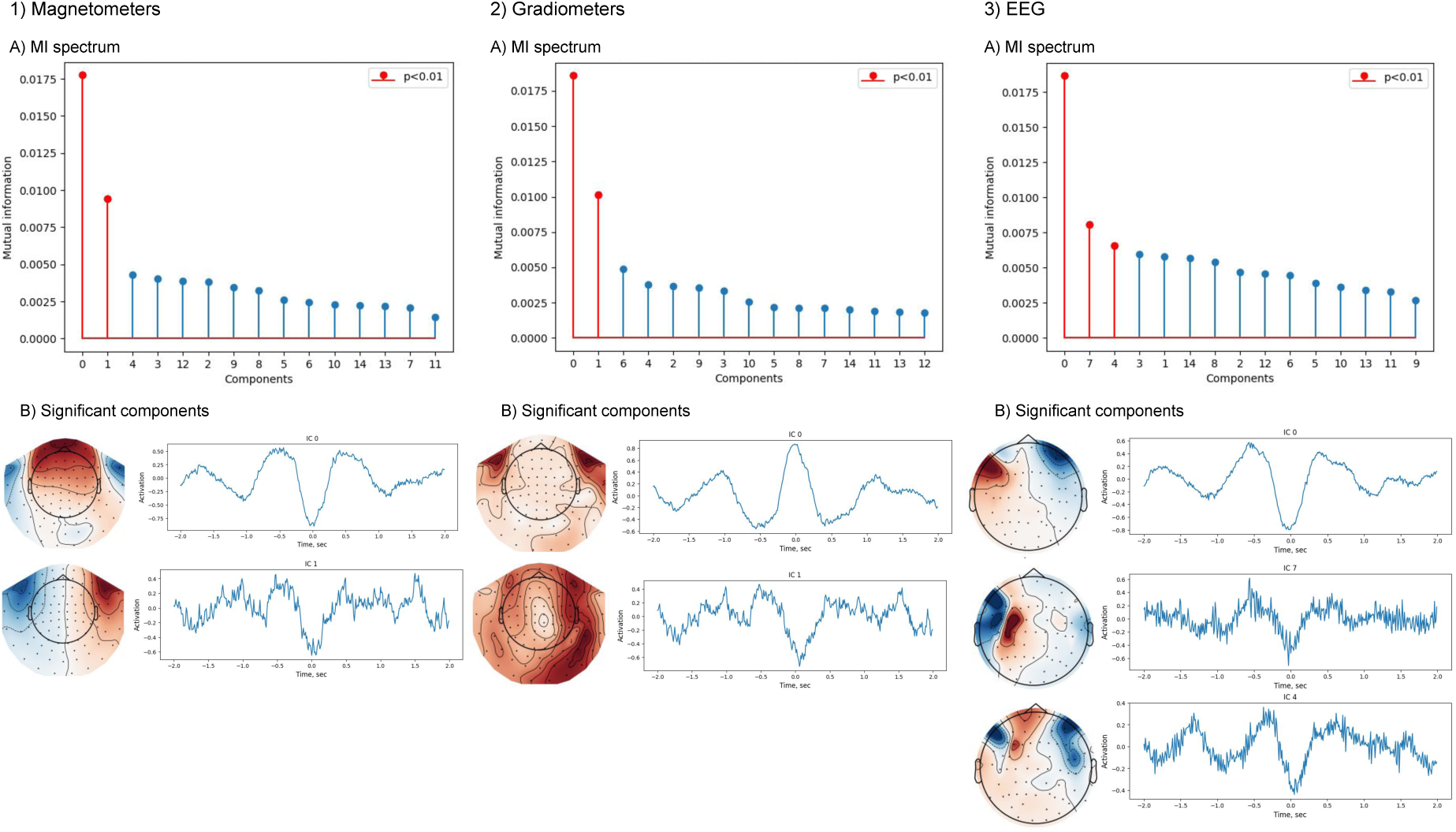
Results of ICA for ingestion artifacts obtained for magnetometers (1), gradiometers (2), and EEG (3): A) MI spectrum illustrating the distribution of mutual information values across ICs. B) Topographies and activation time courses of significant independent components, showing spatial and temporal characteristics of the extracted sources.

These findings highlight the fundamental differences between EEG and MEG in representing both physiological and artifactual signals. For physiological waveforms EEG predominantly captures a smaller number of high-amplitude, spatially diffuse components while MEG decomposes its multivariate waveforms into multiple spatially distinct sources with diverse temporal profiles. This suggests that MEG may provide a more detailed and distributed representation of cortical activity, while EEG remains highly sensitive to dominant, radially oriented sources. For artifacts, the pattern is reversed: EEG extracts a broader range of components, reflecting its susceptibility to sources of non-neuronal origin, whereas MEG selectively isolates a smaller number of components with distinct spatial characteristics.

### SNR estimation

To further evaluate the sensitivity of different sensor modalities to neuronal signals and artifacts, we conducted a detailed analysis of the SNR and the total information capacity for EEG, magnetometer, and gradiometer recordings.

To mitigate overestimation of the total SNR, we performed orthogonalization of the channels, yielding eigenvalues for the matrix of SNR values, see (Iivanainen et al., 2021). The total information, *I*(**A**), was subsequently computed for covariance matrices **A** extracted from the data recorded with three different sensor types. The histograms in Figure 19 illustrate that the total information values obtained from EEG sensors lie to the left of those corresponding to both types of MEG sensors (see Fig. 19).

**Figure 19:**
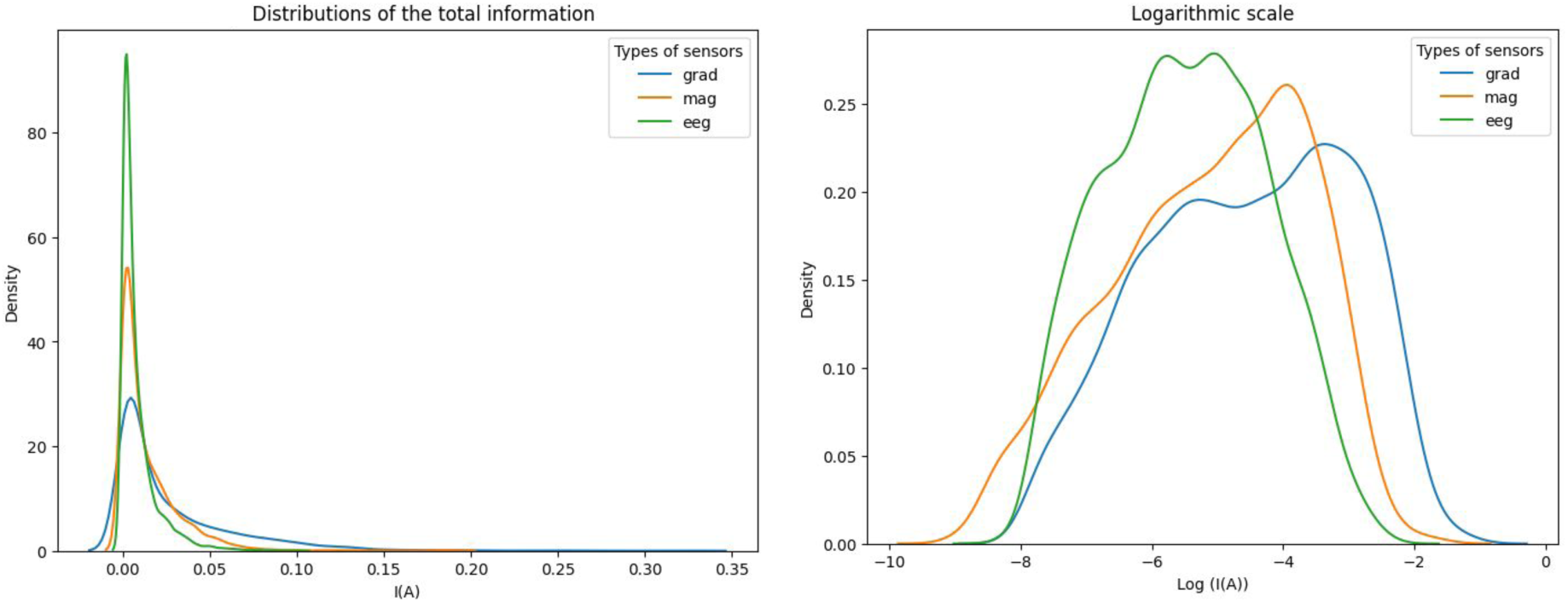
Distribution histograms of total information (I(A)) for EEG, magnetometers, and gradiometers over the linear (left) and logarithmic (right) scales.

The Kruskal-Wallis H-test was conducted to evaluate differences in total information *I*(**A**) across the three sensor types. The test revealed a statistically significant difference among the groups, *H*(2) = 1041.60*, p <* 0.001. Post hoc pairwise comparisons using Dunn’s test with Bonferroni correction confirmed significant differences between all sensor types. Gradiometers (*M* = 0.0272) exhibited significantly higher total information than both magnetometers (*M* = 0.0140*, p <* 0.001) and EEG (*M* = 0.0083*, p <* 0.001). Magnetometers also showed significantly higher total information compared to EEG (*p <* 0.001).

The data-based SNR evaluation was performed using blinks and ingestion as the examples of artifacts and vertex potential and K-complexes as the examples of physiological signals.

The two-way ANOVA was conducted to examine the effects of pattern type (blinks, ingestion, vertex potentials, K-complexes), sensor type (EEG, magnetometers, gradiometers), and their interaction on total information. The analysis revealedsignificant main effects for both pattern type, *F* (3, 2436) = 138.23*, p <* 0.001*, η*^2^ = 0.145, and sensor type, *F* (2, 2436) = 8877.35*, p <* 0.001*, η*^2^ = 0.879. Additionally, a significant interaction was observed between pattern type and sensor type, *F* (6, 2436) = 202.56*, p <* 0.001*, η*^2^ = 0.333.

Post-hoc comparisons using Tukey’s test were conducted to evaluate pairwise differences in total information across the sensor types (EEG, gradiometers, and magnetometers) ((see Fig. 20, A). Gradiometers (*M* = 33.24) demonstrated significantly higher total information compared to EEG (*M* = 13.51*, p <* 0.001). Magnetometers (*M* = 17.66) also exhibited significantly higher total information than EEG (*M* = 13.51*, p <* 0.001). Additionally, gradiometers (*M* = 33.24) showed significantly higher total information compared to magnetometers (*M* = 17.66*, p <* 0.001). These results confirm that gradiometers captured the highest total information among the sensor types, followed by magnetometers, with EEG yielding the lowest values.

**Figure 20:**
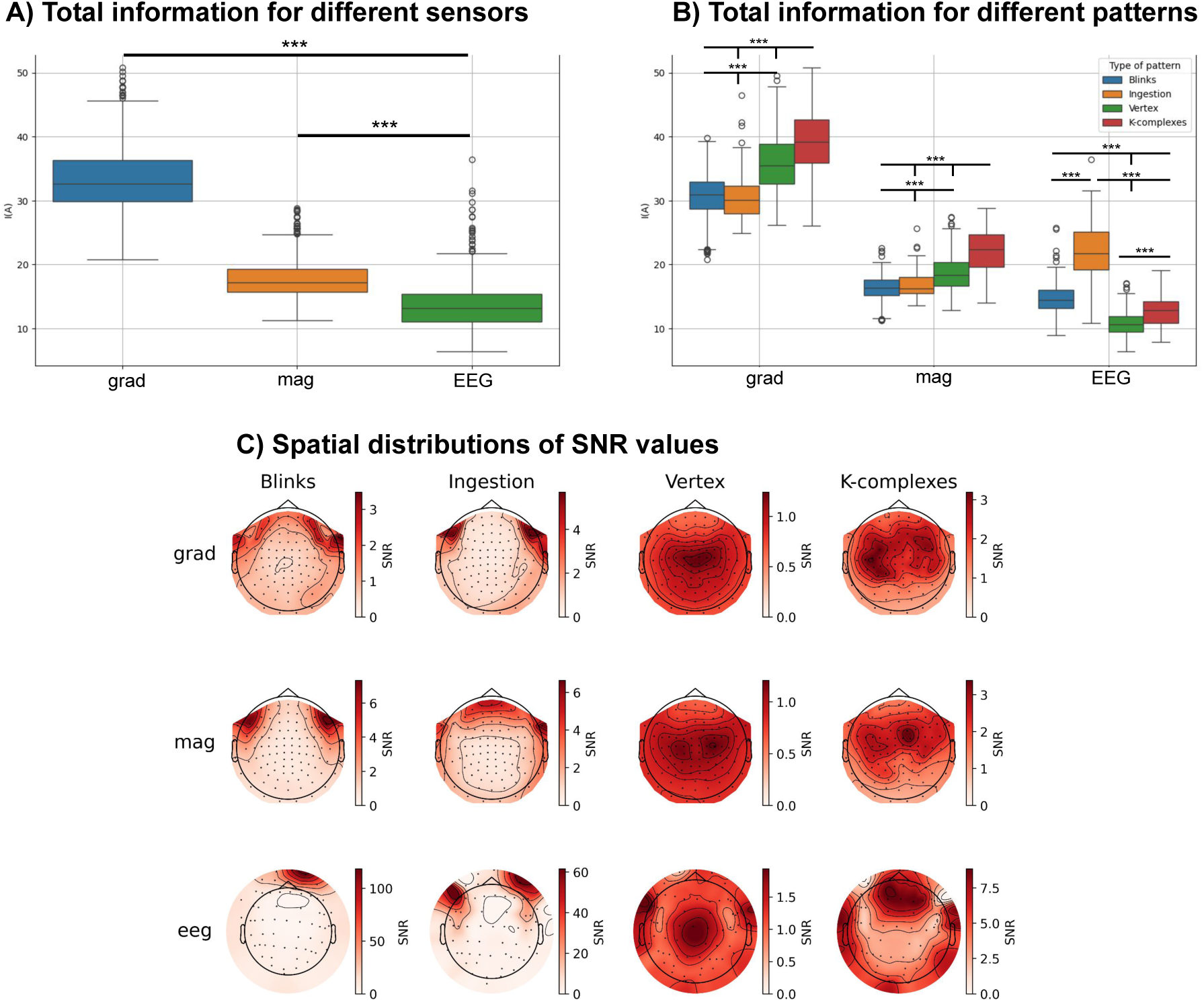
Total information across patterns (blinks, ingestion, vertex potentials, and K-complexes) for each sensor type (gradiometers, magnetometers, and EEG). (A) Comparison between channel types. (B) Comparison between pattern types. (C) Spatial distribution of SNR values.

Additionally, post-hoc comparisons using Tukey’s test were performed to evaluate differences in total information across patterns (blinks, ingestion, vertex potentials, and K-complexes) within each group of channels ((see Fig. 20, B)). For gradiometers, K-complexes (*M* = 39.29) exhibited significantly higher total information compared to blinks (*M* = 30.55*, p <* 0.001), ingestion (*M* = 31.48*, p <* 0.001), and vertex potentials (*M* = 35.82*, p <* 0.001). Vertex potentials (*M* = 35.82) also demonstrated significantly higher total information compared to blinks (*M* = 30.55*, p <* 0.001) and ingestion (*M* = 31.48*, p <* 0.001).

For magnetometers, K-complexes (*M* = 22.12) demonstrated significantly higher total information compared to blinks (*M* = 16.23*, p <* 0.001), ingestion (*M* = 17.18*, p <* 0.001), and vertex potentials (*M* = 18.66*, p <* 0.001). Vertex potentials (*M* = 18.66) also exhibited significantly higher total information compared to blinks (*M* = 16.23*, p <* 0.001) and ingestion (*M* = 17.18*, p <* 0.01).

For EEG, on the contrary, ingestion (*M* = 22.26) exhibited significantly higher total information compared to blinks (*M* = 14.74*, p <* 0.001), K-complexes (*M* = 12.83*, p <* 0.001), and vertex potentials (*M* = 10.73*, p <* 0.001). Blinks (*M* = 14.74) showed significantly higher total information compared to vertex potentials (*M* = 10.73*, p <* 0.001) and K-complexes (*M* = 12.83*, p <* 0.01). Additionally, K-complexes (*M* = 12.83) demonstrated significantly higher total information compared to vertex potentials (*M* = 10.73*, p <* 0.001).

The spatial topographies of SNR values (see Fig. 20, C) closely align with the topographies of the corresponding averaged patterns for each sensor type. This resemblance suggests that the distribution of SNR across sensors reflects the spatial characteristics of the underlying neural and artifact-related signals.

## Discussion

The present study provides a comprehensive qualitative and quantitative comparison of physiological normal variant patterns and artifacts in MEG and EEG. By systematically analyzing common neurophysiological waveforms and non-neuronal interferences, we highlight the complementary nature of these modalities, emphasizing their respective strengths and limitations in capturing brain dynamics. The study was conducted by a team of doctors and neuroscientists, integrating clinical and research perspectives to ensure a multifaceted interpretation of the results.

Our findings reaffirm that MEG and EEG differentially represent physiological activity due to their distinct biophysical properties. The data confirm that EEG is particularly sensitive to radially oriented sources, as evidenced by the clear occipital topography of alpha spindles and the prominent expression of posterior slow waves of youth (PSWoY). At the same time, MEG yields more focal and spatially resolved representations. These differences in MEG and EEG can be explained by heterogeneity of cortical sources underlying the normal variant patterns, and what is observed in EEG as pronounced oscillations actually results from the overlapped divergent cortical activity, which is more accurately disentangled by MEG. Our findings align with previously reported high synchronization of cortical oscillations observed in EEG, as well as the rapid phase, hemispheric, and spatial shifts detected in MEG. (Dehghani et al., 2010). These discrepancies suggest that spindles originate in focal cortical areas (better captured by MEG) before spreading to broader regions detected by EEG, with MEG spindles starting 150 ms earlier and ending 250 ms later than EEG spindles (Dehghani et al., 2011).

However, MEG also presents challenges, as certain patterns may not be visible due to MEG signal strength dependence on source orientation and depth, limiting MEG’s ability to detect certain cortical generators. This underscores the need for developing advanced modeling techniques to enhance interpretation and utility of MEG measurements.

Furthermore, the differential localization of sleep-related waveforms (e.g., sleep spindles, K-complexes, vertex waves) across MEG and EEG underscores their complementary value in sleep research. The localization of vertex waves in our study exhibited a quadripolar pattern extending into sensorimotor regions, suggesting a broader cortical involvement than traditionally described. Prior research dominantly based on MEG has consistently reported that vertex potentials during sleep originate from central brain regions, particularly the sensorimotor cortex. This has been linked to sensory processing during sleep, where vertex sharp transients are associated with primary sensorimotor areas for auditory and tactile stimuli (Stern et al., 2011). Moreover, rhythmic auditory stimulation has been shown to evoke strong vertex responses, closely tied to physiological changes such as cardiac activity(Fruhstorfer et al., 1971). These findings suggest that vertex activity may serve a dual role: facilitating sensory processing during sleep and contributing to motor control despite the absence of voluntary movement. While the exact functional significance of vertex waves remains unclear, one prevailing hypothesis is that they help regulate the arousal threshold in response to sensory stimuli (Stern et al., 2011). Our findings, demonstrating a more distributed activation pattern, may indicate that vertex waves engage a wider cortical network, potentially reflecting a more integrative mechanism underlying their role in sleep-related sensory and motor processes.

MEG recordings in our study showed a frontoparietal dominance of sleep spindles, while EEG reflected a more posterior distribution. This aligns with the thalamic origin of spindles and their projection to different cortical areas via thalamocortical pathways (Fernandez and Lüthi, 2020; Takeuchi et al., 2016). MEG gradiometers, being sensitive to local tangential currents, predominantly capture spindles in frontal and central regions where thalamocortical connectivity is strongest. In contrast, EEG, which detects both tangential and radial dipoles, registers a broader distribution, extending into occipital areas. This is consistent with known spindle propagation patterns, where slow spindles shift anteriorly, while fast spindle activity moves toward posterior regions (Gombos et al., 2022).

Notably, K-complexes localized to the temporal poles in MEG, while in EEG, they were predominantly observed in frontal regions. The physiological interpretation of K- complexes, in particular, requires solving the inverse problem to properly localize and understand their cortical origins, highlighting the methodological challenges involved.

Our results are largely consistent with those reported by Rampp et al. (2020), one of the few studies investigating normal variants in both EEG and MEG. Although the set of examined patterns only partially overlapped (including alpha, SMR (mu), POSTS, vertex waves, and K-complexes), key similarities emerged. Specifically, our findings regarding the parieto-occipital topography of alpha rhythms and the lateralized distribution of mu rhythms align with those described in that paper. Similarly, their characterization of POSTs as an occipital dipole near the midline with a spike-like pattern in MEG corresponds to our localization and observations. However, a notable discrepancy arose in the vertex waves. Rampp et al. (2020) identified midline deep cortical regions as the primary generators of this pattern, which they attributed to the limitations of single equivalent current dipole (ECD) modeling when applied to complex, distributed signals. In contrast, our distributed source modeling approach provided a more comprehensive representation, resolving multiple contributing dipoles in MEG. This suggests that methods relying on distributed rather than single-dipole assumptions may offer superior accuracy in characterizing spatiotemporally complex neurophysiological patterns. This is especially important in cases when the observed data are generated by synchronous sources (Kuznetsova et al., 2021).

In addition to these methodological differences, our study extends beyond the work of Rampp et al. (2020) by incorporating a group-level analysis, averaging across participants and patterns, and performing systematic source modeling for all identified patterns. Moreover, we examined both physiological patterns and artifacts, further enhancing the breadth of our findings.

Physiological and movement-related artifacts were observed in both MEG and EEG, with comparable visual inspection results between the two modalities. Artifacts persisted in MEG even after MaxFilter preprocessing. Although initial visual assessments suggested a similar artifact load in both modalities, deeper quantitative analyses revealed modality-dependent differences in artifact expression.

A critical aspect of our analysis was the evaluation of artifacts that share morphological features with genuine neurophysiological patterns. The ICA results demonstrated that MEG-derived independent components exhibited greater topographical variability, indicative of a more distributed neural origin. Additionally, a larger number of independent components in MEG were associated with physiological activity compared to EEG. This suggests that MEG captures a richer and more complex representation of underlying neural processes, potentially enabling more precise separation of different functional sources. This advantage is particularly relevant for applications requiring fine-grained neurophysiological differentiation, such as epilepsy monitoring and functional connectivity studies. At the same time, the number of artifact-related independent components did not differ significantly between the modalities, indicating that the overall artifact burden remained comparable across MEG and EEG.

The additional SNR analysis revealed significant modality-dependent differences in signal fidelity. The results indicate that MEG gradiometers capture the highest total information, followed by magnetometers and then EEG. However, MEG is also limited by the rapid decay of magnetic field strength with distance, making it less sensitive to deeper cortical sources. The wide dynamic range of patterns detected by MEG highlights its potential but also points to the importance of wearable MEG systems (based on optically pumped magnetometers (OPMs) and YIGM sensors) to improve accessibility and sensitivity to deep sources (Pedersen et al., 2022; Brookes et al., 2022; Razorenova et al., 2024; Koshev et al., 2021).

The total information analysis further demonstrated that MEG provided more distinguishable representations of physiological waveforms than EEG. Notably, vertex potentials and K-complexes exhibited significantly higher total information values in MEG compared to EEG, suggesting that MEG may offer superior precision in detecting transient cortical events. Importantly, total information was higher for physiological patterns in MEG, whereas in EEG, it was greater for artifacts. This suggests that while MEG provides a more detailed representation of neurophysiological activity, EEG remains particularly sensitive to non-neuronal contributions, which can be advantageous for detecting externally induced interferences and assessing their impact on brain signals. Additionally, advancements in amplifier technology suggest that future ultra-high-density EEG systems with reduced noise levels may improve source resolution, potentially bridging the gap in spatial precision between the two modalities (Schreiner et al., 2023, 2024).

The findings presented in this study underscore the necessity of integrating MEG and EEG in neurophysiological research and clinical applications. While EEG remains the gold standard for clinical neurophysiology due to its accessibility and established diagnostic frameworks, MEG offers clear advantages in spatial resolution, artifact rejection, and source localization accuracy. The observed differences between MEG and EEG highlight the potential for multimodal approaches to enhance the detection and characterization of both pathological and physiological brain activity.

For epilepsy research, the improved spatial precision of MEG in identifying cortical generators of epileptiform discharges supports its use as a complementary tool in pre-surgical evaluations. Likewise, the differential sensitivity of MEG and EEG suggests that a combined approach may provide a more comprehensive understanding of neurophysiology and its clinical implications.

## Conclusion

In summary, our results provide strong evidence that MEG and EEG offer distinct yet complementary insights into neurophysiological and artifactual signals. MEG demonstrates superior spatial precision and resilience to the artifacts, while EEG remains invaluable for detecting radially oriented sources and high-amplitude slow-wave activity. However, the challenges associated with MEG interpretation and the limitations in detecting deep sources necessitate ongoing advancements in modeling techniques and hardware innovations such as wearable MEG devices. Given the modality-specific strengths and limitations, future studies should prioritize the integration of both techniques to maximize the accuracy and interpretability of neurophysiological measurements in both research and clinical settings.

## Appendix. SNR estimation

According to (Iivanainen et al., 2021; Petrenko et al., 2021), given the measurement model **x**(*t*) = **G***s*(*t*) + **n**(*t*), where **x**(*t*) is the sensor measurement vector, **G** is the forward model, *s*(*t*) represents the neuronal source activity, and **n**(*t*) is the noise, the channel-wise SNR can be defined as

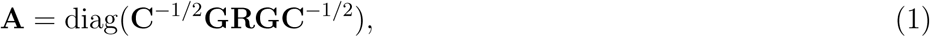

where **C** is the measurement noise covariance and **R** is the covariance of activity of the neuronal sources. Given the pattern at the given epoch *i* **x***_i_*(*t*), the matrix of SNR values **A** was computed as **A** = **C**^−1^*^/^*^2^**YC**^−1^*^/^*^2^, where **Y** is the sensor-space covariance matrix of **x***_i_*(*t*) .

Additionally, in order to avoid overestimation of the total SNR, we performed orthogonalization of the channels and obtained the vector *λ* of eigenvalues of **A**. Then, we used it for the estimation of the total information captured in a single data sample from the set of *M* channels:

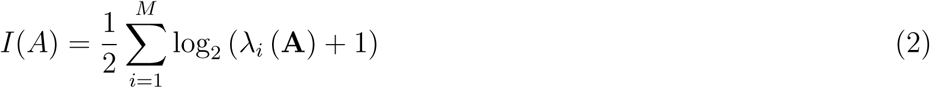

Furthermore, for each neuronal source *i*, under the assumption that only the given source is active, we estimated SNR independent of the observed data :

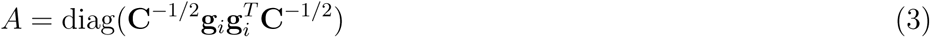

Then, for each source *i* we then calculated the total information as shown in Equation 2 and compared these information maps across the three sensor types: magnetometers, gradiometers, and EEG.

